# Structure and contingency determine mutational hotspots for flower color evolution

**DOI:** 10.1101/2020.08.18.256503

**Authors:** Lucas C. Wheeler, Boswell A. Wing, Stacey D. Smith

## Abstract

Evolutionary genetic studies have uncovered abundant evidence for genomic hotspots of phenotypic evolution, as well as biased patterns of mutations at those loci. However, the theoretical basis for this concentration of particular types of mutations at particular loci remains largely unexplored. In addition, historical contingency is known to play a major role in evolutionary trajectories, but has not been reconciled with the existence of such hotspots. For example, do the appearance of hotspots and the fixation of different types of mutations at those loci depend on the starting state and/or on the nature and direction of selection? Here we use a computational approach to examine these questions, focusing the anthocyanin pigmentation pathway, which has been extensively studied in the context of flower color transitions. We investigate two transitions that are common in nature, the transition from blue to purple pigmentation and from purple to red pigmentation. Both sets of simulated transitions occur with a small number of mutations at just four loci and show strikingly similar peaked shapes of evolutionary trajectories, with the mutations of largest effect occurring early but not first. Nevertheless, the types of mutations (biochemical vs. regulatory) as well as their direction and magnitude are contingent on the particular transition. These simulated color transitions largely mirror findings from natural flower color transitions, which are known to occur via repeated changes at a few hotspot loci. Still, some types of mutations observed in our simulated color evolution are rarely observed in nature, suggesting that pleiotropic effects further limit the trajectories between color phenotypes. Overall, our results indicate that the branching structure of the pathway leads to a predictable concentration of evolutionary change at hotspot loci, but the types of mutations at these loci and their order is contingent on the evolutionary context.

## Introduction

Evolutionary genetic hotspots are the repeated genetic loci of evolution (Martin and Orgogozo 2013; Stem and Orgogozo 2008), appearing across a variety of biological systems. Prominent examples include the roles of *MC1R* in animal melanism, *shavenbaby* in loss of Drosophila trichomes, and several loci of the anthocyanin biosynthesis pathway in floral coloration, all of which have repeatedly experienced mutations underlying phenotypic transitions (Kopp 2009). The repeated involvement of genetic hotspots has been used to argue for the predictability of evolution at the molecular level (Stern and Orgogozo 2008; Streisfeld et al. 2011). Although repeated events are simply a pattern of historical changes, they suggest there is something about hotspot loci that accounts for their over-representation. For example, constraints imposed by the structure of pathways may restrictthe accessibleregion of genotype-phenotypespace, favoring certainpathway targets and restricting the number of possible evolutionary paths (Morrison and Badyaev 2016; Vitkup et al. 2006). Such effects have been observed at the scale of protein evolution, where protein structure and function constrain the order and identity of possible amino acid substitutions (Bridgham et al. 2009; Franzosa and Xia 2009; Harms and Thornton 2014; Weinreich et al. 2006). While we expect similar contingency in the evolution of organismal traits, with constraints on the order of changes at individual loci and across loci involved in phenotypic transitions (Edwards 2019), reconstructing such histories remains significantly more challenging.

Despite the many cases of genomic hotspots underlying phenotypic transitions (Martin and Orgogozo 2013), there are still major gaps in our understanding. First, the appearance of hotspots is likely closely tied to the structure of genetic pathways, but the precise relationship is not well defined. For example, the emergence of a hotspot in a situation where only one genetic mechanism exists is intuitively unsurprising (Shi and Yokoyama 2003), whereas the presence of hotspots in phenotypes that can be achieved via many possible mechanisms is more surprising (Ahnert 2017; Ng and Smith 2016a,b). Second, it remains unclear whether the degree to which the concentration of changes at hotspot loci is due to the intrinsic genetic architecture of the trait or to the external selective forces that manifest as pleiotropic effects (Kopp 2009; Stern and Orgogozo 2008; Streisfeld and Rausher 2011; Wessinger and Rausher 2012). Finally, little is known about the importance of mutational order and the role of context dependence, as observed in protein evolution (Bridgham et al. 2009; Gong et al. 2013; Harms and Thornton 2014; Kent and Green 2017; Salverda et al. 2011; Shah et al. 2015; Starr et al. 2018). For instance, is the involvement of particular loci or types of mutations contingent on the starting state of the system (and thus on changes that occurred before)? Answering these questions will be essential for understanding the basis for repeated targeting of certain loci and the predictability of evolution.

Here we model the evolution of a pathway about which we know a good deal from empirical work, allowing us to make direct comparisons between model predictions and observations from nature. The anthocyanin pathway, which is broadly conserved across flowering plants (Campanella et al. 2014), produces an array of colorful pigments falling into three classes: red pelargonidin-derived pigments, purple cyanidin-derived pigments, and blue delphinidin-derived pigments (Fig. 1A). These pigments are responsible for the most of diversity in coloration across fruits and flowers; they are what make roses red and blueberries blue (Winkel-Shirley 2001). Due to its deeply conserved topology, the anthocyanin pathway has become a prominent system for the study of genetic hotspots in phenotypic evolution (Kopp 2009; Streisfeld and Rausher 2011; Wessinger and Rausher 2012). The pathway is highly branched and reticulated, with multiple instances of competition between enzymes for substrates as well as competition between substrates for enzymes (Fig. 1A).

**Fig 1.**
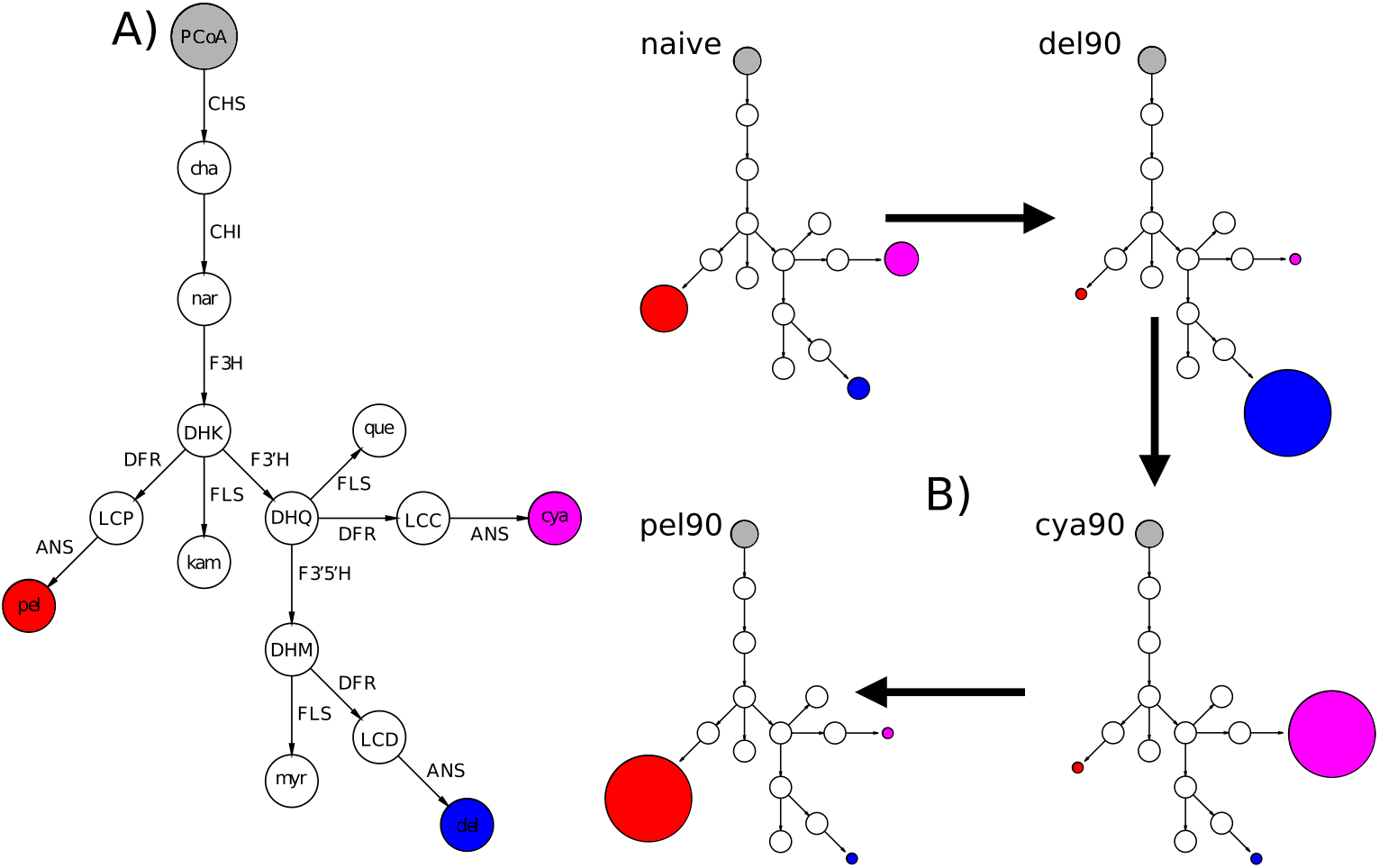
Simulations of color evolution using an anthocyanin pathway model. A) The anthocyanin pathway, a portion of the larger flavonoid biosynthetic pathway (Winkel-Shirley 2001), is shown with substrates at nodes and enzymes along arrows. Flux moves from PCoA at the top down through the branches of the pathway, terminating in the three types of pigments (pelargonidin (pel), cyanidin (cya), and delphinidin (del)), as well as the three flavonols kampferol (kam), quercetin (que), and myricetin (myr). B) The simulations began with a naive state (all pathway parameters equal; see text) and moved first to blue (90% delphinidin), then to purple (90% cyanidin), and finally to red (90% pelargonidin). Substrates include: PCoA (P-Coumaroyl-CoA), cha (chalcone), nar (naringenin), DHK (dihydrokampferol), DHQ (dihydroquercetin), DHM (dihydromyricetin), LCD (leucopelargonidin), LCC (leucocyanidin), LCD (leucodelphinidin). Enzyme abbreviations are: CHS (chalcone synthase), CHI (chalcone isomerase), F3H (flavanone-3-hydroxylase), F3’H (flavonol-3’hydroxylase), F3’5’H (flavonoid-3’5’hydroxylase), DFR (dihyroflavonol-4-reductase), FLS (flavonol synthase), ANS (anthocyanidin synthase).

Taking a computational approach, we explore the range of evolutionary trajectories connecting blue, purple and red phenotypes, and ask how the mutations fixed during these trajectories compare to those observed in nature. In comparing these transitions, we also aim to address general questions about how selected mutations are expected to differ based on the nature of the selection on the pathway. These include (1) Does the location of the phenotypic optimum in pigmentation-space affect the identities of hotspot loci? (2) Are the type, direction, and order of mutations in evolutionary trajectories predictable based on pathway topology and location of the optimum? (3) How does the control of pathway dynamics shift to accommodate new pigment phenotypes? By dissecting a large number of simulated evolutionary trajectories for a sequential set of phenotypic transitions we are able to determine the importance of pathway structure, selective context, and changes to the pathway dynamics in altering the pigmentation phenotype. Our results identify the mechanisms of pathway evolution, generate testable predictions for future empirical studies, and provide a generally-applicable framework of the emergence and behavior of hotspot loci based on interactions between pathway topology and selection.

## Methods

### Design of the kinetic model of the anthocyanin pathway

We constructed a kinetic model of the anthocyanin biosynthetic pathway that is designed to capture the most salient pathway features. The model consists of a set of differential equations (described below; also detailed in the Supplemental text) that capture the topology and dynamics of a simplified representation of the anthocyanin pathway. It includes the branches that produce the anthocyanidin pigments (pelargonidin, cyanidin, and delphinidin), as well as the competing set of branches that produce the flavonols (kampferol, quercetin, and myricetin) from shared precursor compounds (Fig. 1A; Fig. S1). We incorporated several simplifying assumptions into the model, such as ignoring the linked production of flavone compounds (Winkel-Shirley 2001) and the decision to exclude the overlap in activity between the F3’H and F3’5’H enzymes, because this shared activity is variable in nature (Falginella et al. 2010; Kaltenbach et al. 1999; Seitz et al. 2007).

To represent the pathway reactions, we used a generalized Michaelis-Menten rate law formulation (Chou and Talaly 1977). We specified irreversible rate laws for each enzymatic reaction in the pathway model using the Tellurium library (Choi et al. 2018), as previously described (see Supplemental text and (Wheeler and Smith 2019)). Each enzyme rate law in the pathway has three different types of parameters: *K*_*cat*_; the catalytic turnover rate, *K*_*m*_; the Michaelis constant (related to a dissociation constant), and *E*_*t*_; the enzyme concentration. This rate law formulation allows us to incorporate substrate competition for enzymes with multiple substrates, such as DFR, by using unique *K*_*cat*_ and *K*_*m*_ parameters for the different substrates (see Fig. 1A). We previously determined that irreversible rate laws were a reasonable approximation for the behavior of the anthocyanin pathway (Wheeler and Smith 2019) and a simple irreversible model has also been shown to fit well to the kinetic results of *in vivo* experiments. An added advantage is that the irreversible model requires fewer parameters. To allow calculation of a steady state solution for the system of equations that represent the pathway dynamics, we instituted two boundary processes: 1) the incoming upstream concentration of PCoA (Fig. 1) is fixed at a constant value, 2) the rate of transport of the products out of the pathway system is determined by *K*_*sink*_ (a rate constant shared by all final products of the pathway: pelargonidin, cyaninidin, delphinidin, kampferol, quercetin, and myricetin; see Fig. 1) multiplied by the product concentration (see model specification in e.g. Supplemental File “cyanidin-to-pelargonidin-simulations.py”).

### Design scheme for the evolutionary simulations

We simulated sequential phenotypic transitions (Fig. 1B) using the evolutionary algorithm implemented in our python package enzo (Wheeler and Smith 2019) (https://github.com/lcwheeler/enzo). Briefly, our algorithm works by sampling “mutations” from a gamma distribution with *α* = 0.8 and *β* = 3. These mutations are multiplicative shifts that can result in either an increase or decrease to the value of a single *K*_*cat*_, *K*_*m*_, or *E*_*t*_ parameter. Each iteration of the algorithm introduces a random mutation to a randomly selected parameter and then re-calculates the steady state concentration of all chemical species in the pathway. Since the focus of this study is on shifts among pigments (as opposed to changes in the amount of pigmentation), we discarded mutations that altered total steady state production beyond a 10% tolerance. This experimental design is in line with empirical work, where species can shift between pigment types while keeping the total anthocyanin levels relatively constant (Berardi et al. 2016; Esfeld et al. 2018), resulting in flowers of different hues but similar color intensity. Natural and engineered systems also evidence trade-offs between anthocyanins and flavonols, suggesting constraints on total flavonoid content (Davies et al. 2003; Nakatsuka et al. 2007; Nielsen et al. 2002; Yuan et al. 2016). For mutations falling within the tolerance, a selection coefficient is calculated using the fitness function: *W* = exp(−(*ratio*_*current*_*−ratio*_*opt*_)^2^), which depends on the distance of the steady state ratio of a target pigment to the sum of all pigments (*ratio*_*current*_) from a pre-defined phenotypic optimum (*ratio*_*opt*_) (Clark 1991; Rausher 2013; Wheeler and Smith 2019; Wright and Rausher 2010). A fixation probability is calculated based on a formula (1 − *e*^*−s*^) that is weighted by the selection coefficient. The mutation is then either fixed or discarded probabilistically. Neutral (*s* = 0) and deleterious (*s <* 0) mutations are discarded for efficiency, because their fixation probabilities are negligible. The algorithm performs a series of iterations until either the optimum is reached within a defined tolerance (here 10%) or a pre-set maximum number of iterations (here 50,000) is reached.

Previously, we used this framework to study the transition from a “naive” pathway model (all kinetic parameters of a certain type are initialized with equal numerical values) to a state wherein 90% of the total steady state concentration of all chemical species was composed of the blue delphinidin pigment (see (Wheeler and Smith 2019)). Here, we focused on transitions between naturally occuring phenotypes (blue, purple and red flowers) (Fig. 1B). The initial starting state was the original naive pathway model mentioned above. We evolved the model from the naive phenotype to a state wherein delphinidin composed 90% of the total *anthocyanin* concentration at steady state (the “blue” phenotype), with a total of 10,000 unique simulated trajectories. We then calculated the mean evolved value for each model parameter and initialized a new starting state with these values. We evolved this mean 90% delphinidin state (hereafter referred to as simply the delphinidin state) to a 90% cyanidin state (corresponding to “purple”; hereafter referred to as simply the cyanidin state) and finally repeated the procedure to simulate the transition from cyanidin to 90% pelargonidin (corresponding to “red”; hereafter referred to as the pelargonidin state). Organizing the simulations around this stepwise sequence of transitions allowed us to circumvent issues that arise from epistasis in the pathway model, wherein different starting states can result in differences between trajectories that are difficult to interpret (Ng et al. 2018).

Finally, we implemented two additional simulation experiments to examine the effect of the branched pathway structure on the properties of evolutionary trajectories. First, we created a simplified linear pathway model (the sub-pathway containing only the steps leading to pelargonidin production, initialized with the parameters from the naive model), and evolved this model toward an optimum defined by a 3-fold increase in steady-state pelargonidin concentration relative to the starting state. Second, we imposed a constraint that the ratio of pelargonidin at steady state to all other pathway substrates/products remain constant. To examine the effect of the fitness function alone, we then applied this same scenario the (branched) naive anthocyanin pathway model. We carried out 2,000 replicates of each of these simulations. See Supplemental methods for details on the subsequent analyses of all simulations.

## Results

### Evolution of hotspot enzymes is dependent on the phenotypic transition

Although the two trajectories were of similar length and overlapped in the loci targeted, we observed marked differences in the type and frequency of mutations at pathway enzymes. The median length for trajectories in both phenotypic transitions is four steps (see trajectory length distributions in Fig. S2). Roughly 99% of all fixed mutations in both transitions are spread across four loci, F3’5’H, F3’H, DFR, and FLS (Fig. S3), which we identified as pathway hotspots in our earlier study (Wheeler and Smith 2019). However, the frequencies of fixation at these hotspot enzymes depend on the phenotypic transition under selection (Fig. 2A). The predominant switch is in the fixation frequencies of mutations to F3’5’H and F3’H. In the transition from blue delphinidin to purple cyanidin, mutations at F3’H are fixed much more frequently than those at F3’5’H. This pattern is reversed in the transition from purple cyanidin to red pelargonidin, where fixed mutations to F3’5’H parameters are relatively rare (Fig. 2A). In comparison, the fixation frequency of mutations at DFR and FLS are quite similar in both transitions. These results demonstrate the dependence of hotspot behavior on the axis of variation under selection.

**Fig 2.**
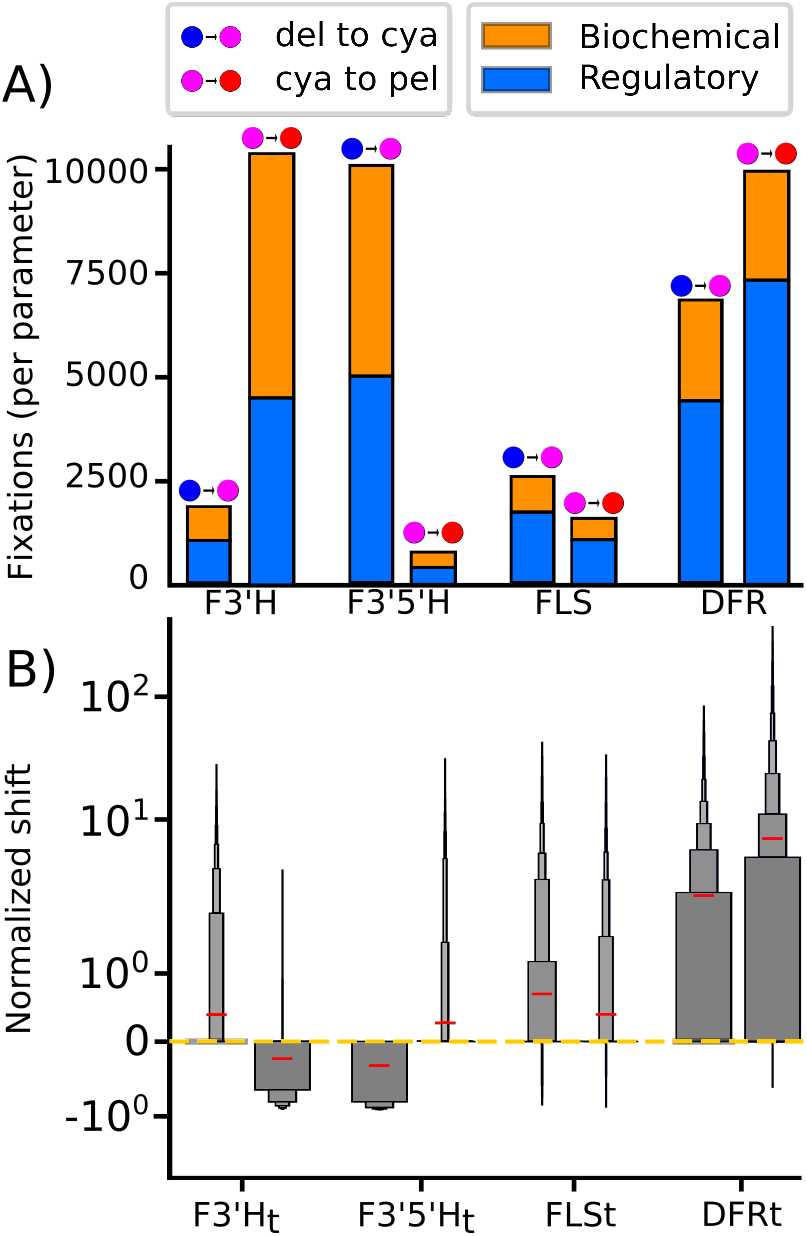
Hotspots vary depending on transition type with consistent patterns of mutation. Comparison of fixed mutations the four hotspot loci. The delphinidin to cyanidin and cyanidin to pelargonidin transitions are labeled (see Fig. 1B). A) Overall proportions of biochemical and regulatory mutations for each hotspot locus in both phenotypic transitions, normalized by the total number of each parameter type per enzyme (e.g. DFR has three *K*_*cat*_ values and three *K*_*M*_ values for a total of six “biochemical” parameters). Stacked orange and blue boxes show the proportion of biochemical (*K*_*M*_ and *K*_*cat*_) and regulatory (*E*_*t*_) mutations (see legend). B) Boxenplot (Hofmann et al. 2017) of normalized directional shifts for the concentration parameter from each of the four enzymes shown in panel A. Box width is proportional to the amount of data in enclosed region. The mean of each distribution is shown by horizontal red line. Directional shifts are shown on a log scale. Distributions of directional shifts for the *K*_*M*_, *K*_*cat*_, and *E*_*t*_ for all the enzymes in the pathway are shown in Fig. S6.

We found that there is an overall bias toward regulatory mutations (enzyme concentration), compared with biochemical mutations (changes to kinetic parameters), in both phenotypic transitions. Regulatory mutations represent *>* 50% of all fixations at the four hotspot enzymes. Nonetheless, both biochemical and regulatory mutations are highly represented in the trajectories of both transitions (Fig. 2A). Changes in concentration of F3’H and F3’5’H can easily drive shifts in the relative production of downstream products due to their location at branch points that commit flux toward one or another pigment branch (Fig. 1A). However, the reason for the effects of regulatory changes at DFR and FLS is less immediately obvious. We previously observed that the naive pathway has an inherent bias toward pelargonidin/kampferol production, due simply to the early partitioning of flux down these committed branches. Thus, a shift in DFR concentration affects the steady state production along each branch differently (Fig. S4). These differences can be accentuated or suppressed by differential biochemical changes to DFR activity on pigment precursors (see Supplemental text and Fig. S5 for a detailed explanation of this general phenomenon). Likewise, the ability of FLS to draw flux away from all anthocyanin pigment branches allows it to redirect flux to tune the anthocyanin output in a manner that is also dependent on the relative efficiency of the competing reactions.

We found that the phenotypic transitions are characterized by predictable sets of positive and negative mutations at hotspot enzymes. Specifically, certain parameters at the hotspot loci are shifted in a way that increases a particular reaction, while others reduce the activity of a particular reaction. The distributions of these shifts for the concentration parameters are shown in Fig. 2B (see Fig. S6 for the shift distributions of all *E*_*t*_, *K*_*cat*_, and *K*_*M*_ parameters). As with the fixation frequencies at each enzyme, these directional changes are also contingent on which phenotypic transition is being made (Fig. 2B, Fig. S6). The most striking example can be seen in the shift distributions for concentration parameters of F3’H and F3’5’H in Fig. 2B. In the transition from the delphinidin to the cyanidin, fixed mutations to F3’H concentration are almost exclusively positive, while mutations to F3’5’H concentration are negative. This pattern is exactly reversed in the transition from cyanidin to pelargonidin (Fig. 2B).

### Trajectories have consistent structure with biased order of fixed mutations

Looking across simulations, the curves for fixed mutations show a skewed distribution, with the largest mutations occurring in the early (but not earliest) steps (Fig. 3). There is a peak around the second trajectory step representing the largest selection coefficients (Fig. 3; note: this is despite the aggregate distributions of selection coefficients for all fixed mutations regardless of trajectory step being roughly exponentially-distributed, see Fig. S7). We confirmed that this peaked pattern in the stepwise distributions was not an artifact of aggregating data from trajectories, by ranking the selection coefficients of fixed mutations in each individual trajectory. In both the blue to purple and the purple to red transitions, we observed that in *>* 50% of trajectories the selection coefficient of the mutation fixed in the second step was larger than that in the first step. This observation demonstrates that typical trajectories exhibit a pattern similar to that shown by the aggregated distribution curves in Fig. 3.

**Fig 3.**
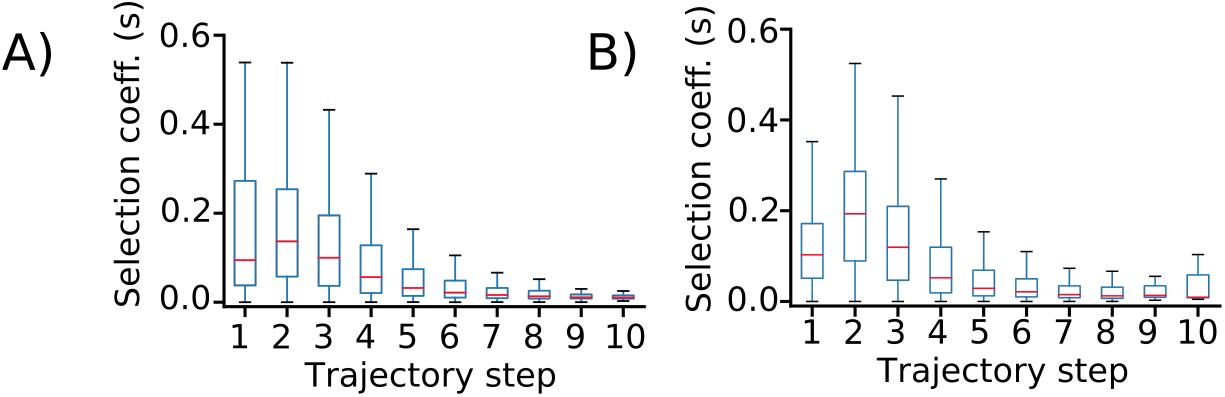
Largest fixed mutations occur at early (but not earliest) trajectory steps. Distributions of *fixed* selection coefficients at each trajectory step over all 10,000 simulations per phenotypic transition, shown as boxplots for A) delphinidin to cyanidin and B) cyanidin to pelargonidin. The curves are peaked around the second step, resembling a chemical activation barrier. Red lines indicate median values.

We hypothesized that the peaked shape of the selection coefficient curves could be due to the interplay between our chosen fitness function and the complex interactions within the pathway. These interactions could restrict the availability of largest-effect mutations at the beginning of trajectories. As described in the Methods section, we tested this idea by employing two additional simulation experiments. A simple linear pathway model (containing only the series of reactions leading to pelargonidin), evolved under selection for increased absolute pelargonidin production with fixed ratio of pelargonidin to the other species (see Methods), exhibited a roughly exponential distribution of selection coefficients over trajectory steps (Fig. S8). This result suggests that the branching structure of our main model could contribute to the peaked stepwise distributions of selection coefficients during adaptive walks. However, when we applied the same fitness function to the naive pathway model (with the full branching structure), we observed a similar exponential distribution of selection coefficients (Fig. S9). This result indicates that the combination of directional selection on target-pigment ratio and stabilizing selection on total pathway production (see Methods) is likely the main cause of the peaked shape of the adaptive walks. It is worth noting that the fitness function used in these additional simulations (see Methods) results in increased frequency of fixed mutations at both upstream and downstream pathway enzymes (Fig. S10). This observation is consistent with the imposed selection on overall increased concentration of an end-product (requiring increased total pathway flux) rather than selection to maintain the total flux of the starting state.

We next examined the series of mutations underlying the two focal transitions. Despite relying on mutations at different parameters, both the blue to purple and purple to red transitions showed a similar tight relationship between the sensitivity of the pathway to particular mutations and the position of those mutations in the trajectories. Specifically, mutations at enzyme parameters with higher mean sensitivity (see Methods) were biased toward earlier fixation in trajectories, as can be seen by the similarity in heat maps in Fig. 4. The result of higher sensitivity is that, for those parameters, a small mutation to the parameter value can induce a bigger shift in pathway function on average. Since there is a general trend for larger-effect mutations to be fixed earlier in trajectories, these sensitivity differences manifest in certain parameters becoming concentrated among early mutations while others are skewed toward the middle or end of trajectories (Fig. 4). There is also a positive relationship between sensitivity and total number of fixed mutations at a given parameter (Fig. 4). We also noted that the parameters contributing to the cyanidin to pelargonidin transition is almost entirely a subset of those that contribute to the delphinidin to cyanidin transition, a pattern resulting from the effective clipping of the delphinidin branch during the first simulated transition. Nonetheless, the order of mutations at those parameters differs markedly between trajectories (dashed lines in Fig. 4), indicative of the corresponding differences in sensitivities. For example, the blue delphinidin state is highly sensitive to mutations in k_DFR_DHQ (the *K*_*cat*_ for DFR on the DHQ precursor), making it an early target, while the purple cyanidin state has little sensitivity to mutations in that parameter, leaving them for later in the trajectories.

**Fig 4.**
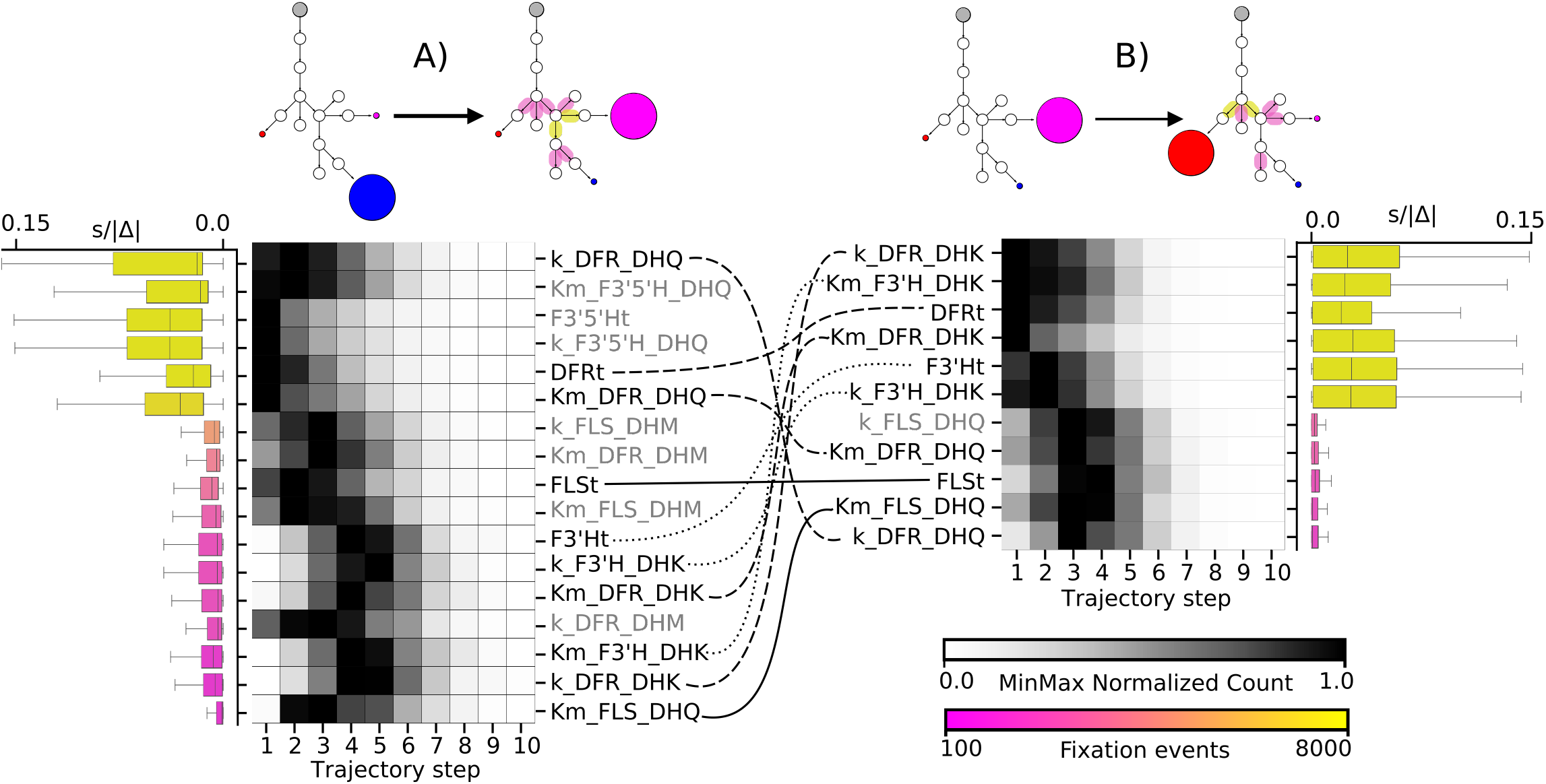
Order of fixed mutations strongly influenced by sensitivity. For each transition (delphinidin to cyanidin in A and cyanidin to pelargonidin in B), heatmaps show the distribution of fixation events across the trajectory steps for all parameters making up at least 1% of all fixed mutations. Parameter names are listed in descending order of total number of fixation events (colored pink to yellow). Boxes in the heatmap show the fixation events at each step in the trajectories across the 10,000 simulations; counts are normalized on a MinMax (0,1) scale for comparison (see MinMax Normalized Count scale bar). Boxplots show the *complete* distributions of “sensitivity” values (|*s/*Δ|; absolute value of the ratio of selection coefficient (s) the normalized directional mutation size (Δ): (*mutant value − previous value*)/(*previous value*)) for *all* mutations for the given parameter (see Fig. S11 for explanation of the sensitivity calculation). Reactions involving the more-frequently-targeted/higher-sensitivity parameters (yellow) and less-frequently-targeted/lower-sensitivity parameters (pink) are highlighted on the pathway diagrams at the top of figure. Parameter notation: *K*_*m*_ = *K*_*M*_ (e.g. Km_DFR_DHQ), *k* = *K*_*cat*_ (e.g. k_DFR_DHQ), and *Et* = *E*_*t*_ (e.g. DFRt), where the substrate associated with the kinetic parameters is indicated (e.g. k_DFR_DHQ). Parameters shared across transitions are connected by curved lines, where line style is matched for parameters of the same enzyme: DFR (dashed), F3’H (dotted), and FLS (solid). Parameters unique to each transition are written in gray.

### Control of pathway dynamics is re-arranged to achieve switches between pigment phenotypes

The primary change to the pathway dynamics during each phenotypic transition is a re-arrangement of the underlying structure of control that enzymes exert over steady state concentrations of chemical species in the pathway (see Methods). The predominant shift in the pathway control structure is summarized in Fig. 5A. The pathway is effectively linearized down the path (series of reactions) leading to the target pigment, e.g. cyanidin or pelargonidin. This re-arrangement transforms the path to the target pigment into a path of reduced resistance. These shifts in pathway control are largely reproducible, falling within a fairly narrow range across the majority of simulations (Fig. 5A, Fig. S13).

**Fig 5.**
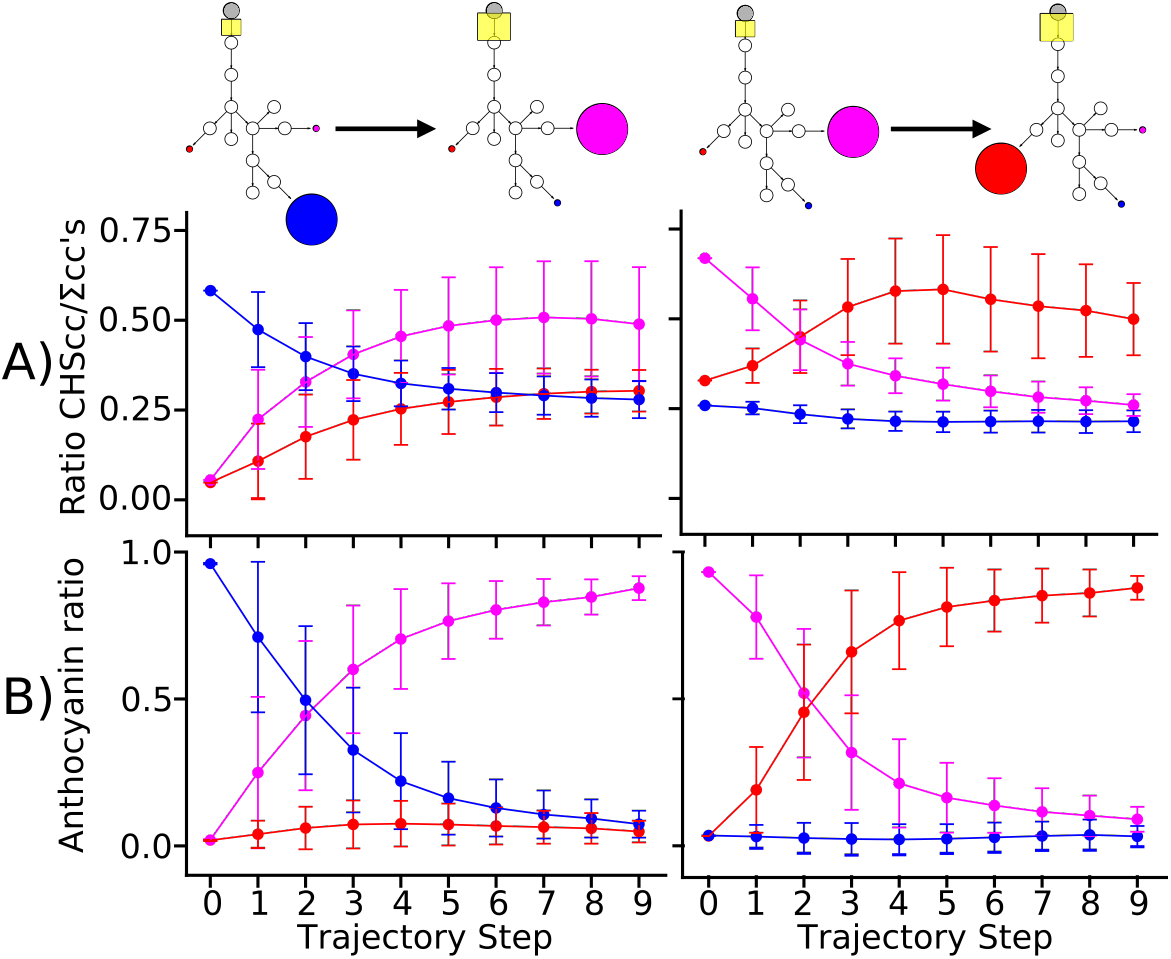
Pathway flux dynamics are re-arranged to accommodate phenotypic transitions. Left column is delphinidin to cyanidin transition, right column is cyanidin to pelargonidin transition (see pathway diagrams at the top). A) Trajectories through the concentration control coefficient (CC) space, depicting the absolute value of the ratio of the CC of the CHS enzyme for each pigment to the sum of all CC values for that species 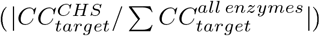 across all trajectory steps. The position of CHS in the pathway is highlighted by the yellow boxes in pathway diagrams at the top, with the change in size denoting increased relative control over target pigment. The proportion of total control over the target species possessed by CHS increases to saturation in both cases. Inflection points of these curves align with inflection points of the pigment ratio curves in panel B. The overshoot in the relative control of CHS over pelargonidin (red) during the transition from cyanidin (purple) to pelargonidin is likely due to the intrinsic bias of the path-way toward pelargonidin production. Error bars are one standard deviation. B) Mean trajectories through pigment space (in relative amounts of each); the relative amount of the target pigment is proportional to fitness in our model. Colors follow above (*delphinidin* = *blue*, *cyanidin* = *purple*, *pelargonidin* = *red*). Error bars are the standard deviation of the mean.

The linearization effect is achieved by weakening the control that the branching enzymes (such as F3’5’H and F3’H; see Fig. S12) have over the target pigment, while strengthening the control they have over the off-target species. Particularly for the other two non-target anthocyanin pigments, the branching enzyme control coefficients become more negative, which can be interpreted as an stronger dampening of off-target pigment production (Fig. S12). The net effect of these shift is to grant the dominant control over flux down the target path (i.e. cyanidin or pelargonidin) to CHS, the furthest upstream enzyme in the pathway, which is itself negligible as a direct target of fixed mutations (Fig. 1A, Fig. 5A, Fig. S12). The trajectories of the absolute value of the ratio of the CHS concentration control coefficient for the target pigment to the sum of all coefficients for that pigment are shown in Fig. 5A. In both transitions this ratio converges to a stationary value over the course of simulations, mirroring the trajectories through pigment space shown in Fig. 5B.

The mean trajectories of the target pigment ratio (relative to the total anthocyanins produced at steady state) are the phenotype-level manifestation of the underlying changes to the pathway control structure. They follow a sigmoidal curve across trajectory steps (Fig. 5B). Shifts in flux down the target branch result in trade-offs with other branches that are particularly pronounced between the current target species and the target species from the previous transition in the sequence. This is due to the constraint on the total amount of material produced by the pathway, wherein as some pathway products increase, the material that is allotted to others decreases. The state-dependency of these trade-offs demonstrates the emergence of epistasis from the simple kinetic model.

## Discussion

### Color transitions occur via changes at a subset of pathway loci

In simulations of pathway evolution, we largely recover the same hotspot loci as seen in nature. For example, transitions from blue delphinidin to red pelargonidin production in *Iochroma* and *Penstemon* (Smith and Rausher 2011; Smith et al. 2013; Wessinger and Rausher 2014, 2015), blue delphinidin to purple cyanidin in *Phlox* (Hopkins and Rausher 2011), and purple cyanidin to red pelargonidin in *Ipomoea* (Marais and Rausher 2010) were accomplished by mutations at F3’5’H, F3’H, and DFR. In fact, these three loci, particularly F3’H and F3’5’H, are consistently the causative targets of fixed mutations underlying color variation in nature (reviewed in (Wheeler and Smith 2019); see also Table S1). Our simulations clearly demonstrate that the hotspot nature of these enzymes is derived from their position in the pathway structure, which imparts a relatively higher ability to control flux. In other words, these are the loci that are able to perturb the pigmentation phenotype sufficiently to be selected on. This observation is in line with empirical findings in other metabolic pathways, such as those responsible for carotenoid biosynthesis in plants, wherein pathway position strongly affects the rate of fixed mutations at pathway loci (Clotault et al. 2012). The overall effect of this narrowing of mutational targets is to impart some predictability and repeatability to evolution of the pathway, particularly at the scale of which enzymes are likely to be involved. However, the dependence of mutational effects on the selected phenotypic transition demonstrates the need for knowledge of the precise context in which the pathway is evolving in order to make accurate predictions. For example, in cases of parallel evolution of a new pigmentation phenotype from a known common ancestral state, it will likely be possible to predict the targets and types of mutations (Wessinger and Rausher 2012).

Further consistent with our simulated results, natural transitions in anthocyanin pigmentation are often accomplished with a small number of functional changes, although these may be composed of several mutational steps at the genetic level. For example, the blue-to-red transition in *Iochroma* was achieved by the combined effect of three changes (deactivation of F3’5’H, down-regulation of F3’h, and coding mutations in DFR) (Smith and Rausher 2011), while the *Phlox* transition required two (coding mutations at *F3’5’h* and a Myb transcription factor that regulates *F3’5’h* expression). Meanwhile, the Penstemon transition required only one change (deactivation of F3’5’H) as did the *Ipomea* transition (down-regulation of *F3’h*) (Marais and Rausher 2010). Similarly small numbers of fixed changes have also been observed in a variety of other evolutionary transitions, such as changes in floral morphology in *Mimulus* (Bradshaw et al. 1998), selection for growth rate in *Aspergillus nidulans* (Schoustra et al. 2009), adaptation to nutrient-limitation in *Saccharomyces cerevisiae*, and adaptive melanism in *Chaetodipus* mice (Nachman et al. 2003; Steiner et al. 2007). This pattern reinforces the notion that major transitions can have a relatively simple genetic basis and can thus happen on short timescales given a sufficiently high rate for the generation of genetic variation (Gervasi and Schiestl 2017).

One notable difference between the set of hotspot enzymes so far known in nature and that of our simulations is the flavonol synthase (FLS) enzyme, a common target of fixed mutations in the pathway model. FLS is the enzyme responsible for the synthesis of the flavonol compounds (kampferol, quercetin, and myricetin) from the DHK, DHQ, and DHM precursors that are shared with the anthocyanin branches (Fig. 1). The flavonols play a variety of critical biological roles including acting as sunscreen compounds that shield tissues from UV radiation (Ryan et al. 1998, 2002b). Our modeling shows that mutations at FLS can alter anthocyanin pigmentation by increasing activity on the shared precursors and thus drawing flux away from the anthocyanin branches. Changes in FLS concentration can yield differential effects depending on the status of the other enzymes catalyzing reactions that lead down any particular anthocyanin path. This result highlights the fact that, because of interactions among loci in the pathway, even those enzymes not required for anthocyanin biosynthesis can affect pigmentation. To our knowledge, only one empirical study of flower color variation has identified FLS as a target. In this case, differential regulation of FLS induces flower color pattern variation by tuning anthocyanin concentration rather than causing a shift between pigments (Yuan et al. 2016). However, it is worth noting that many searches for color loci have focused on candidate genes required for anthocyanin biosynthesis, so FLS has not received much attention. Additionally, FLS has been targeted to induce a switch between two anthocyanin pigments in genetically-engineered systems (Table S1) (Nakatsuka et al. 2007; Nielsen et al. 2002). Based on our simulated results, FLS can therefore be thought of as a hotspot for “missing mutations” that we may expect to observe often in nature, but so far have only observed very rarely.

Another pathway enzyme that is rarely, if ever, involved in color transitions in nature is ANS (Table S1). In contrast to FLS, this observation is easily rationalized by our model. ANS has very little differential control over flux down the anthocyanin branches, because it is at the very end of the pathway (Fig. 1; Fig. S12). By the time precursors reach ANS, the bulk of relative flux partitioning down the pathway branches has already been determined by the upstream F3’H, F3’5’H, and DFR; that partitioning is then relatively insensitive to changes at ANS. Nevertheless, ANS was a more common target of fixed mutations when selecting on increased absolute concentration in the simple linear pathway control (see Methods; Fig. S10), consistent with the ability of ANS to potentially act as a final “bottleneck” for overall flux of pathway material toward anthocyanin production. These observations are in line with the occasional involvement of ANS as a target in transitions between pigmented and un-pigmented phenotypes in flowers (Keiichi Shimizu 2011) and fruits (Rafique et al. 2016).

### Range of targets points to pleiotropic effects of biochemical mutations in nature

While we largely recovered the same hotspot loci as those observed in nature, the types of mutations in our simulations are broader. Specifically, although regulatory mutations (represented by enzyme concentration in our model) were preferred at all four hotspot loci, we also observed a substantial fraction of fixed biochemical mutations (those changing kinetic enzyme parameters). This contrasts with what is so far known about pigmentation transitions in nature, wherein changes at some loci are predominantly regulatory while changes at others tend to be structural (Wessinger and Rausher 2012). *F3’h*, in particular, has been primarily targeted by regulatory mutations in nature (Table S1), although it has also occasionally been targeted by biochemical activity-tuning modifications (Hoshino et al. 2003; Zufall and Rausher 2003). In contrast, direct modifications of enzyme activity (in particular the alteration of specificity for anthocyanidin precursors) appear to be much more common than regulatory mutations at DFR (Table S1). This may be due to the need for modifications that alter DFR activity to tune pathway flux after shifts induced by regulatory mutations at branching enzymes, which redirect flux down the target branch (Smith and Rausher 2011; Smith et al. 2013; Zufall and Rausher 2004).

This discrepancy in the distribution of mutation types between our model and empirical observations may be explained by pleiotropic effects that occur in nature, particularly for mutations that affect enzyme function. Most of the enzymes in the anthocyanin pathway play multiple roles yet are encoded by single-copy-number genes (Winkel-Shirley 2001). Therefore, changes in the enzymatic activity at the protein level would affect the enzymes in all tissues where they are expressed. If the change in activity induced by the mutation is detrimental to other functions when occurring in some tissues, this would impose a constraint on the allowable mutations (Wagner and Zhang 2011). In contrast, a tissue-specific regulatory mutation could avoid this antagonistic pleiotropy by allowing activity in the off-target tissue to remain unchanged (Streisfeld and Rausher 2011; Streisfeld et al. 2011; Wessinger and Rausher 2012). As an example, in the case of simulated transitions from cyanidin production to pelargonidin production, we observe changes which worsen effectiveness of F3’H on DHQ, the precursor for both leucocyanidin and quercetin. This will have the side effect of lowering quercetin production, and since quercetin is a more effective sunscreen than the monohydroxylated kaempferol produced from DHK (Ryan et al. 1998, 2002a), we would expect negative fitness effects from these changes. Alternatively, expression of *F3’h* is known to be under the control of different transcription factors in different tissues (Smith and Rausher 2011; Streisfeld and Rausher 2009), setting up the potential availability of tissue-specific regulatory mutations that can alter the function while avoiding antagonistic pleiotropy. In contrast to *F3’h*, *F3’h5’h* is thought to be a less pleiotropic gene overall, consistent with its propensity to undergo complete, system-wide loss-of-function mutations (Wessinger and Rausher 2012) (Table S1).

Although we believe that pleiotropy explains many of the differences between our model and nature, aspects of our simulation design may also play a role. Our simulations treat the distribution of parameter mutations equally for all pathway parameters, because the natural distributions are not known. It is possible that the relative probability of encountering an advantageous mutation of a certain type differs across pathway loci due to biophysical constraints. At DFR for example, there are many examples of mutations at different positions in the enzyme sequence that can yield functional changes (reviewed in (Smith et al. 2013)). At other enzymes, there may be fewer sites that can readily achieve the necessary functional effects due to biophysical limitations arising from the structure or chemistry of the enzyme (Copley 2009; Kaltenbach and Tokuriki 2014; Khersonsky and Tawfik 2010). Thus, even if the mutation rate is the same across enzymes, there may still be less raw variation for selection to act upon. Future empirical work to measure the effects of spontaneous mutations, and the way they are shaped by the structural and kinetic properties of these enzymes in nature, would allow us to better model the available variation in individual enzyme genes.

### Evolutionary walks between phenotypes avoid largest steps first

Our simulations suggest that although color shifts can occur with a few mutations, those with largest effect (in terms of selection coefficient) do not tend to get fixed first. With a median of 4 selected mutations to reach the target phenotype, the largest mutations are typically fixed at the second step. This runs counter to some well-established theoretical results for adaptive walks, which predict an exponential distribution of mutation sizes over trajectory steps (Joyce et al. 2008; Orr 1998, 2002; Rokyta et al. 2005). However, it is worth noting that these studies have several limitations including the assumption that all the mutations are drawn from the same unchanging distribution of fitness effects and involve a single locus (Joyce et al. 2008; Orr 1998). We suspect that the deviation of our model from the theoretical prediction may be due in part to the violation of the typical assumptions, in particular that the branched pathway contains complex interactions between loci. As described in the Results, evolution of a simplified linear pathway did in fact yield a roughly exponential distribution of fixed selection coefficients. However, it appears that the constraints imposed by the fitness function used in our model (directional toward a specific target-pigment ratio optimum and stabilizing on total steady state concentration) had a larger effect on the distribution. We argue that, under this fitness function, very large-effect mutations that move the pathway phenotype in the correct direction are difficult to achieve, and thus rare, at the beginning of trajectories. In contrast, it appears that moderately-sized mutations can disrupt the system sufficiently to open up new possible paths for subsequent mutations to facilitate further movement toward the optimum.

One of the key observations of this study is that the concept of “large effect” vs. “small effect” mutations is not fixed for a change of a given size at a given parameter. The effect size of any given parameter change on the pathway fitness is instead contingent on the location of the phenotypic optimum and the distance of the pathway phenotype from the optimum at each step of a trajectory. Thus the distribution of mutational effects on fitness shifts as the pathway evolves, rather than remaining static. We suggest that these dynamics are likely to be common for phenotypes controlled by intertwined networks of genes or enzymes, where epistasis is expected to be pervasive. Interestingly, a similar adaptive curve to those observed in this study (sigmoidal in the fitness space) emerges from a simple biophysical model of transcription-factor network evolution (Lässig 2007) and peaked distributions can also be observed in models of evolution with shifting optima (Collins et al. 2007). Furthermore, under certain extreme value domains fitness effects are expected to increase, rather than decrease, over the course of adaptive walks (Seetharaman and Jain 2014).

It is difficult to determine how well the observed order of effect sizes for fixed changes align with nature. Studies of the evolution of polygenic phenotypes in plants that have been done, using approaches such as QTL mapping, yield all of the differences between evolutionary end states, but the order in which they accumulated remains unknown (Bradshaw et al. 1998; Hodges et al. 2002). Furthermore, the effects of these mapped regions are likely to be different in any extant proxy organism than they would have been in the ancestral organism due to evolutionary separation and the accompanying effects of genetic background (Leips and Mackay 2000; Lunzer et al. 2010). Nonetheless, we do expect mutations with a range of effects on the phenotype, some large and some smaller. In fact, this has been observed in at least one natural flower color transition; blue to red flowers in *Iochroma* (where the change at *F3’h* has the largest effect, followed by that at *F3’h5’h*, and finally by the change at DFR) (Smith and Rausher 2011). Future studies could potentially employ a phylogenetic approach, coupling ancestral state reconstruction with measurements of mutational effect sizes in species with the ancestral pigmentation phenotype. This approach could be used to infer the order of mutations and the stepwise distribution of effect sizes, but great care would need to be taken to account for epistasis between mutations that have occurred along different lineages (Lunzer et al. 2010; Mackay 2014).

## Conclusions

Although the anthocyanin pathway is just one metabolic pathway, the features that it possesses, such as branches and points of substrate competition, are commonplace in metabolism. We therefore expect that the evolutionary dynamics of hotspot loci observed in our simulations will be generally applicable across a wide variety of pathways. Our simulations give insight into how interactions among pathway genes shape movements between phenotypes. First, we found that the starting and ending states have a large effect on evolutionary trajectories, including the loci of evolution, the types of mutations and the order in which they are fixed. These results underscore that “hotspots” are not defined by the pathway alone, but are contingent on the evolutionary context. We expect this context-dependence to be a universal property of complex networks and may be a major contributor to the determination of hotspot loci for any phenotypic transition in nature. Thus, further studying the interplay between pathway position and direction of selection will be critical to understand the repeatability and predictability of evolution. For pathways like the anthocyanin pathway, which are deployed in different tissues, at different times and under different environmental conditions, pleiotropic effects of mutations are likely to further constrain accessible pathways between phenotypes. Computational models, like this one, allow us to determine the types of mutations likely to contribute to phenotypic evolution due to the structure of the pathway alone. ‘Missing’ mutations, those observed in simulations but not in natural systems, point to important gaps in our understanding of natural variation and how selection acts upon it.

## Acknowledgments

We thank members of the Smith lab for their comments on development of the manuscript. We thank Mike Harms and his group at University of Oregon for helpful suggestions regarding analysis of the simulated data. In particular we thank Joseph Harman for insightful comments on the figures. We also thank Zach Sailer of the Jupyter project and Dan Weinreich of Brown University, for their questions and comments that helped to hone the interpretation of our results. This work utilized the RMACC Summit supercomputer, which is supported by the National Science Foundation (awards ACI-1532235 and ACI-1532236), the University of Colorado Boulder, and Colorado State University. The Summit supercomputer is a joint effort of the University of Colorado Boulder and Colorado State University.

## Funding

This work was funded by a RIO Innovative Seed Grant Program (RISGP) award from the University of Colorado-Boulder (LCW) and by NSF-DEB 1553114 (SDS). The funders had no role in study design, data collection and analysis, decision to publish, or preparation of the manuscript.

## Availability of data and materials

The python library enzo) for running evolutionary simulations, is available on github (https://github.com/lcwheeler/enzo). The scripts used to run the simulations, Jupyter notebooks and scripts for the subsequent analyses, and the raw simulated datasets are available in an OSF online repository (https://osf.io/kxr23/). Additional information is available in the supplemental text.

## Author contributions

LCW and SDS conceived the study and outlined the computational approach. LCW constructed the mathematical model, wrote the code, performed the simulations, conducted the data analysis, and generated figures. LCW, BAW, and SDS collaborated on writing the manuscript. SDS, LCW, and BAW secured funding for the work. All authors have read and approved the manuscript.

## Competing interests

The authors declare that they have no competing interests.

## Notes

https://osf.io/kxr23/

### Competing Interest Statement

The authors have declared no competing interest.

## References

Ahnert, S. E. Structural properties of genotype–phenotype maps. Journal of The Royal Society Interface, 14(132):20170275, 2017. doi: 10.1098/rsif.2017.0275.

Berardi, A. E., Hildreth, S. B., Helm, R. F., Winkel, B. S., and Smith, S. D. Evolutionary correlations in flavonoid production across flowers and leaves in the iochrominae (solanaceae). Phytochemistry, 130:119–127, 2016. ISSN 0031-9422. doi: https://doi.org/10.1016/j.phytochem.2016.05.007.

Bradshaw, H. D., Otto, K. G., Frewen, B. E., McKay, J. K., and Schemske, D. W. Quantitative trait loci affecting differences in floral morphology between two species of monkeyflower (mimulus). Genetics, 149(1):367–382, 1998. ISSN 0016-6731.

Bridgham, J. T., Ortlund, E. A., and Thornton, J. W. An epistatic ratchet constrains the direction of glucocorticoid receptor evolution. Nature, 461(7263):515–519, Sep 2009. ISSN 1476-4687. doi: 10.1038/nature08249.

Campanella, J. J., Smalley, J. V., and Dempsey, M. E. A phylogenetic examination of the primary anthocyanin production pathway of the plantae. Botanical Studies, 55(1):10, Jan 2014. ISSN 1999-3110. doi: 10.1186/1999-3110-55-10.

Choi, K., Medley, J. K., König, M., Stocking, K., Smith, L., Gu, S., and Sauro, H. M. Tellurium: An extensible python-based modeling environment for systems and synthetic biology. Biosystems, 171:74–79, 2018. ISSN 0303-2647. doi: https://doi.org/10.1016/j.biosystems.2018.07.006.

Chou, T. C. and Talaly, P. A simple generalized equation for the analysis of multiple inhibitions of michaelis-menten kinetic systems. J. Biol. Chem., 252(18):6438–6442, Sept. 1977. ISSN 0021-9258, 1083-351X.

Clark, A. G. Mutation-selection balance and metabolic control theory. Genetics, 129(3):909–923, Nov. 1991. ISSN 0016-6731, 1943-2631.

Clotault, J., Peltier, D., Souffiet-Freslon, V., Briard, M., and Geoffriau, E. Differential selection on carotenoid biosynthesis genes as a function of gene position in the metabolic pathway: A study on the carrot and dicots. PLOS ONE, 7(6):1–13, 06 2012. doi: 10.1371/journal.pone.0038724.

Collins, S., de Meaux, J., and Acquisti, C. Adaptive walks toward a moving optimum. Genetics, 176(2):1089–1099, 2007. ISSN 0016-6731. doi: 10.1534/genetics.107.072926.

Copley, S. D. Evolution of efficient pathways for degradation of anthropogenic chemicals. Nature Chemical Biology, 5(8):559–566, Aug 2009. ISSN 1552-4469. doi: 10.1038/nchembio.197.

Davies, K. M., Schwinn, K. E., Deroles, S. C., Manson, D. G., Lewis, D. H., Bloor, S. J., and Bradley, J. M. Enhancing anthocyanin production by altering competition for substrate between flavonol synthase and dihydroflavonol 4-reductase. Euphytica, 131(3):259–268, Jun 2003. ISSN 1573-5060. doi: 10.1023/A:1024018729349.

Edwards, E. J. Evolutionary trajectories, accessibility and other metaphors: the case of c4 and cam photosynthesis. New Phytologist, 223(4):1742–1755, 2019. doi: 10.1111/nph.15851.

Esfeld, K., Berardi, A. E., Moser, M., Bossolini, E., Freitas, L., and Kuhlemeier, C. Pseudogenization and resurrection of a speciation gene. Curr. Biol., 28(23):3776–3786.e7, Dec. 2018. ISSN 0960-9822.

Falginella, L., Castellarin, S. D., Testolin, R., Gambetta, G. A., Morgante, M., and Di Gaspero, G. Expansion and subfunctionalisation of flavonoid 3’,5’-hydroxylases in the grapevine lineage. BMC Genomics, 11(1):562, Oct 2010. ISSN 1471-2164. doi: 10.1186/1471-2164-11-562.

Franzosa, E. A. and Xia, Y. Structural Determinants of Protein Evolution Are Context-Sensitive at the Residue Level. Molecular Biology and Evolution, 26(10):2387–2395, 07 2009. ISSN 0737-4038. doi: 10.1093/molbev/msp146.

Gervasi, D. D. L. and Schiestl, F. P. Real-time divergent evolution in plants driven by pollinators. Nature Communications, 8(1):14691, Mar 2017. ISSN 2041-1723. doi: 10.1038/ncomms14691.

Gong, L. I., Suchard, M. A., and Bloom, J. D. Stability-mediated epistasis constrains the evolution of an influenza protein. eLife, 2:e00631, may 2013. ISSN 2050-084X. doi: 10.7554/eLife.00631.

Harms, M. J. and Thornton, J. W. Historical contingency and its biophysical basis in glucocorticoid receptor evolution. Nature, 512(7513):203–207, Aug 2014. ISSN 1476-4687. doi: 10.1038/nature13410.

Hodges, S. A., Whittall, J. B., Fulton, M., and Yang, J. Y. Genetics of floral traits influencing reproductive isolation between aquilegia formosa and aquilegia pubescens. . The American Naturalist, 159(S3):S51–S60, 2002. doi: 10.1086/338372. PMID: 18707369.

Hofmann, H., Wickham, H., and Kafadar, K. Letter-value plots: Boxplots for large data. Journal of Computational and Graphical Statistics, 26(3):469–477, 2017. doi: 10.1080/10618600.2017.1305277.

Hopkins, R. and Rausher, M. D. Identification of two genes causing reinforcement in the texas wildflower *Phlox* drummondii. Nature, 469(7330):411–414, Jan. 2011. ISSN 1476-4687.

Hoshino, A., Morita, Y., Choi, J.-D., Saito, N., Toki, K., Tanaka, Y., and Iida, S. Spontaneous mutations of the flavonoid 3’-hydroxylase gene conferring reddish flowers in the three morning glory species. Plant Cell Physiol, 44(10):990–1001, Oct. 2003. ISSN 0032-0781.

Joyce, P., Rokyta, D. R., Beisel, C. J., and Orr, H. A. A general extreme value theory model for the adaptation of dna sequences under strong selection and weak mutation. Genetics, 180(3): 1627–1643, 2008. ISSN 0016-6731. doi: 10.1534/genetics.108.088716.

Kaltenbach, M. and Tokuriki, N. Dynamics and constraints of enzyme evolution. J. Exp. Zool. (Mol. Dev. Evol.), 322(7):468–487, Nov. 2014. ISSN 1552-5015.

Kaltenbach, M., Schröder, G., Schmelzer, E., Lutz, V., and Schröder, J. Flavonoid hydroxylase from catharanthus roseus: cdna, heterologous expression, enzyme properties and cell-type specific expression in plants. The Plant Journal, 19(2):183–193, 1999. doi: 10.1046/j.1365-313X.1999.00524.x.

Keiichi Shimizu, Nanako Onishi, N. M. A. I. S. O. L. U. Y. S. F. H. A 94-bp deletion of anthocyanidin synthase gene in acyanic flower lines of lisianthus eustoma grandiflorum (raf.) shinn.]. Journal of the Japanese Society for Horticultural Science, 80(4):434–442, 2011. doi: 10.2503/jjshs1.80.434.

Kent, D. G. and Green, A. R. Order matters: The order of somatic mutations influences cancer evolution. Cold Spring Harbor Perspectives in Medicine, 7(4), 2017. doi: 10.1101/cshperspect.a027060.

Khersonsky, O. and Tawfik, D. S. Enzyme promiscuity: A mechanistic and evolutionary perspective. Annu. Rev. Biochem., 79(1):471–505, 2010.

Kopp, A. Metamodels and phylogenetic replication: A systematic approach to the evolution of developmental pathways. Evolution, 63(11):2771–2789, 2009. doi: 10.1111/j.1558-5646.2009.00761.x.

Lässig, M. From biophysics to evolutionary genetics: statistical aspects of gene regulation. BMC Bioinformatics, 8(6):S7, Sep 2007. ISSN 1471-2105. doi: 10.1186/1471-2105-8-S6-S7.

Leips, J. and Mackay, T. F. C. Quantitative trait loci for life span in drosophila melanogaster: Interactions with genetic background and larval density. Genetics, 155(4):1773–1788, 2000. ISSN 0016-6731.

Lunzer, M., Golding, G. B., and Dean, A. M. Pervasive cryptic epistasis in molecular evolution. PLOS Genetics, 6(10):1–10, 10 2010. doi: 10.1371/journal.pgen.1001162. URL https://doi.org/10.1371/journal.pgen.1001162.

Mackay, T. F. C. Epistasis and quantitative traits: using model organisms to study gene-gene interactions. Nature Reviews Genetics, 15(1):22–33, Jan 2014. ISSN 1471-0064. doi: 10.1038/nrg3627.

Marais, D. L. D. and Rausher, M. D. Parallel evolution at multiple levels in the origin of humming-bird pollinated flowers in ipomoea. Evolution, 64(7):2044–2054, 2010. ISSN 1558-5646.

Martin, A. and Orgogozo, V. The loci of repeated evolution: A catalog of genetic hotspots of phenotypic variation. Evolution, 67(5):1235–1250, May 2013. ISSN 1558-5646.

Morrison, E. S. and Badyaev, A. V. Structuring evolution: biochemical networks and metabolic diversification in birds. BMC Evol. Biol., 16(1):168, Aug. 2016. ISSN 1471-2148.

Nachman, M. W., Hoekstra, H. E., and D’Agostino, S. L. The genetic basis of adaptive melanism in pocket mice. Proceedings of the National Academy of Sciences, 100(9):5268–5273, 2003. ISSN 0027-8424. doi: 10.1073/pnas.0431157100.

Nakatsuka, T., Abe, Y., Kakizaki, Y., Yamamura, S., and Nishihara, M. Production of red-flowered plants by genetic engineering of multiple flavonoid biosynthetic genes. Plant Cell Rep, 26(11): 1951–1959, Nov. 2007. ISSN 1432-203X.

Ng, J. and Smith, S. D. How to make a red flower: the combinatorial effect of pigments. AoB PLANTS, 8, Jan. 2016a.

Ng, J. and Smith, S. D. Widespread flower color convergence in solanaceae via alternate biochemical pathways. New Phytol., 209(1):407–417, 2016b. ISSN 1469-8137.

Ng, J., Freitas, L. B., and Smith, S. D. Stepwise evolution of floral pigmentation predicted by biochemical pathway structure. Evolution, 72(12):2792–2802, 2018. ISSN 1558-5646.

Nielsen, K., Deroles, S. C., Markham, K. R., Bradley, M. J., Podivinsky, E., and Manson, D. Anti-sense flavonol synthase alters copigmentation and flower color in lisianthus. Molecular Breeding, 9(4):217–229, Dec 2002. ISSN 1572-9788. doi: 10.1023/A:1020320809654.

Orr, H. A. The population genetics of adaptation: The distribution of factors fixed during adaptive evolution. Evolution, 52(4):935–949, 1998. doi: 10.1111/j.1558-5646.1998.tb01823.x.

Orr, H. A. The population genetics of adaptation: The adaptation of dna sequences. Evolution, 56 (7):1317–1330, 2002. doi: 10.1111/j.0014-3820.2002.tb01446.x.

Rafique, M. Z., Carvalho, E., Stracke, R., Palmieri, L., Herrera, L., Feller, A., Malnoy, M., and Martens, S. Nonsense mutation inside anthocyanidin synthase gene controls pigmentation in yellow raspberry (rubus idaeus l.). Frontiers in Plant Science, 7:1892, 2016. ISSN 1664-462X. doi: 10.3389/fpls.2016.01892.

Rausher, M. D. The evolution of genes in branched metabolic pathways: Evolution in branched pathways. Evolution, 67(1):34–48, Jan. 2013. ISSN 00143820.

Rokyta, D. R., Joyce, P., Caudle, S. B., and Wichman, H. A. An empirical test of the mutational landscape model of adaptation using a single-stranded dna virus. Nature Genetics, 37(4):441–444, Apr 2005. ISSN 1546-1718. doi: 10.1038/ng1535.

Ryan, K. G., Markham, K. R., Bloor, S. J., Bradley, J. M., Mitchell, K. A., and Jordan, B. R. Uvb radiation induced increase in quercetin: Kaempferol ratio in wild-type and transgenic lines of petunia. Photochem. Photobiol., 68(3):323–330, 1998. ISSN 1751-1097.

Ryan, K. G., Swinny, E. E., Markham, K. R., and Winefield, C. Flavonoid gene expression and uv photoprotection in transgenic and mutant petunia leaves. Phytochemistry, 59(1):23–32, 2002a. ISSN 0031-9422. doi: https://doi.org/10.1016/S0031-9422(01)00404-6.

Ryan, K. G., Swinny, E. E., Markham, K. R., and Winefield, C. Flavonoid gene expression and uv photoprotection in transgenic and mutant petunia leaves. Phytochemistry, 59(1):23–32, Jan. 2002b. ISSN 0031-9422.

Salverda, M. L. M., Dellus, E., Gorter, F. A., Debets, A. J. M., van der Oost, J., Hoekstra, R. F., Tawfik, D. S., and de Visser, J. A. G. M. Initial mutations direct alternative pathways of protein evolution. PLOS Genetics, 7(3):1–11, 03 2011. doi: 10.1371/journal.pgen.1001321.

Schoustra, S. E., Bataillon, T., Gifford, D. R., and Kassen, R. The properties of adaptive walks in evolving populations of fungus. PLOS Biology, 7(11):1–10, 11 2009. doi: 10.1371/journal.pbio.1000250. URL https://doi.org/10.1371/journal.pbio.1000250.

Seetharaman, S. and Jain, K. Adaptive walks and distribution of beneficial fitness effects. Evolution, 68(4):965–975, 2014. doi: 10.1111/evo.12327.

Seitz, C., Ameres, S., and Forkmann, G. Identification of the molecular basis for the functional difference between flavonoid 3’-hydroxylase and flavonoid 3’,5’-hydroxylase. FEBS Letters, 581 (18):3429–3434, 2007. doi: 10.1016/j.febslet.2007.06.045.

Shah, P., McCandlish, D. M., and Plotkin, J. B. Contingency and entrenchment in protein evolution under purifying selection. Proceedings of the National Academy of Sciences, 112(25):E3226–E3235, 2015. ISSN 0027-8424. doi: 10.1073/pnas.1412933112.

Shi, Y. and Yokoyama, S. Molecular analysis of the evolutionary significance of ultraviolet vision in vertebrates. Proceedings of the National Academy of Sciences, 100(14):8308–8313, 2003. ISSN 0027-8424. doi: 10.1073/pnas.1532535100.

Smith, S. D. and Rausher, M. D. Gene loss and parallel evolution contribute to species difference in flower color. Mol. Biol. Evol., 28(10):2799–2810, Oct. 2011. ISSN 0737-4038.

Smith, S. D., Wang, S., and Rausher, M. D. Functional evolution of an anthocyanin pathway enzyme during a flower color transition. Mol. Biol. Evol., 30(3):602–612, Mar. 2013. ISSN 0737-4038.

Starr, T. N., Flynn, J. M., Mishra, P., Bolon, D. N. A., and Thornton, J. W. Pervasive contingency and entrenchment in a billion years of hsp90 evolution. Proceedings of the National Academy of Sciences, 115(17):4453–4458, 2018. ISSN 0027-8424. doi: 10.1073/pnas.1718133115.

Steiner, C. C., Weber, J. N., and Hoekstra, H. E. Adaptive variation in beach mice produced by two interacting pigmentation genes. PLOS Biology, 5(9):1–10, 08 2007. doi: 10.1371/journal.pbio.0050219.

Stern, D. L. and Orgogozo, V. The loci of evolution: How predictable is genetic evolution? Evolution, 62(9):2155–2177, 2008. doi: 10.1111/j.1558-5646.2008.00450.x.

Streisfeld, M. A. and Rausher, M. D. Genetic changes contributing to the parallel evolution of red floral pigmentation among ipomoea species. New Phytol., 183(3):751–763, 2009. ISSN 1469-8137.

Streisfeld, M. A. and Rausher, M. D. Population genetics, pleiotropy, and the preferential fixation of mutations during adaptive evolution. Evolution, 65(3):629–642, 2011. ISSN 1558-5646.

Streisfeld, M. A., Liu, D., and Rausher, M. D. Predictable patterns of constraint among anthocyanin-regulating transcription factors in ipomoea. New Phytol., 191(1):264–274, July 2011. ISSN 0028646X.

Vitkup, D., Kharchenko, P., and Wagner, A. Influence of metabolic network structure and function on enzyme evolution. Genome Biology, 7(5):R39, May 2006. ISSN 1474-760X. doi: 10.1186/gb-2006-7-5-r39.

Wagner, G. P. and Zhang, J. The pleiotropic structure of the genotype-phenotype map: the evolvability of complex organisms. Nature Reviews Genetics, 12(3):204–213, Mar 2011. ISSN 1471-0064. doi: 10.1038/nrg2949.

Weinreich, D. M., Delaney, N. F., DePristo, M. A., and Hartl, D. L. Darwinian evolution can follow only very few mutational paths to fitter proteins. Science, 312(5770):111–114, 2006. ISSN 0036-8075. doi: 10.1126/science.1123539.

Wessinger, C. A. and Rausher, M. D. Lessons from flower colour evolution on targets of selection. J. Exp. Bot., 63(16):5741–5749, Oct. 2012. ISSN 0022-0957.

Wessinger, C. A. and Rausher, M. D. Predictability and irreversibility of genetic changes associated with flower color evolution in penstemon barbatus. Evolution, 68(4):1058–1070, 2014. ISSN 1558-5646.

Wessinger, C. A. and Rausher, M. D. Ecological transition predictably associated with gene degeneration. Mol. Biol. Evol., 32(2):347–354, Feb. 2015. ISSN 0737-4038.

Wheeler, L. C. and Smith, S. D. Computational Modeling of Anthocyanin Pathway Evolution: Biases, Hotspots, and Trade-offs. Integrative and Comparative Biology, 59(3):585–598, 05 2019. ISSN 1540-7063. doi: 10.1093/icb/icz049.

Winkel-Shirley, B. Flavonoid biosynthesis. a colorful model for genetics, biochemistry, cell biology, and biotechnology. Plant Physiol., 126(2):485–493, June 2001. ISSN 0032-0889.

Wright, K. M. and Rausher, M. D. The evolution of control and distribution of adaptive mutations in a metabolic pathway. Genetics, 184(2):483–502, Feb. 2010. ISSN 0016-6731, 1943-2631.

Yuan, Y.-W., Rebocho, A. B., Sagawa, J. M., Stanley, L. E., and Bradshaw, H. D. Competition between anthocyanin and flavonol biosynthesis produces spatial pattern variation of floral pigments between mimulus species. PNAS, 113(9):2448–2453, Mar. 2016. ISSN 0027-8424, 1091-6490.

Zufall, R. A. and Rausher, M. D. The Genetic Basis of a Flower Color Polymorphism in the Common Morning Glory (Ipomoea purpurea). Journal of Heredity, 94(6):442–448, 11 2003. ISSN 0022-1503. doi: 10.1093/jhered/esg098.

Zufall, R. A. and Rausher, M. D. Genetic changes associated with floral adaptation restrict future evolutionary potential. Nature, 428(6985):847–850, Apr. 2004. ISSN 1476-4687.

