## Supplemental text for "Structure and contingency determine mutational hotspots for flower color evolution"

### Structure and contingency determine mutational hotspots for flower color evolution: Supplemental text and figures

### Rate laws for the pathway model

In the equations below the parameters are denoted as follows: $K_{cat}$ parameters indicate the catalytic constant, written with the enzyme name in the superscript and the substrate name in the subscript (for example $K^{CHS}_{cat,PCoA}$). $K_M$ parameters indicate the Michaelis constant, written with the substrate name in the superscript and the enzyme name in the subscript (for example $K^{PCoA}_{M,CHS}$). $v$ indicates velocity of the reaction, written with the enzyme name in the superscript and the product name in the subscript (for example $v^{CHS}_{cha}$). Floating species abbreviations are: PCoA (P-Coumaroyl-CoA), cha (chalcone), nar (naringenin), DHK (dihydrokampferol), DHQ (dihydroquercetin), DHM (dihydromyricetin), que (quercetin), kam (kampferol), myr (myricetin), LCD (leucopelargonidin), LCC (leucocyanidin), LCD (leucodelphinidin), pel (pelargonidin), cya (cyanidin), del (delphinidin). Del, cya, and pel are the anthocyanidins, which are glycosylated to form the various anthocyanins. Kam, que, and myr are the flavonols. Enzyme abbreviations are: CHS (chalcone synthase), CHI (chalcone isomerase), F3H (flavanone-3-hydroxylase), F3'H (flavonol-3'hydroxylase), F3'5'H (flavonoid-3'5'hydroxylase), DFR (dihyroflavonol-4-reductase), FLS (flavonol synthase), ANS (anthocyanidin synthase) (see Fig. S1).

---

##### CHS

$$v^{CHS}_{cha} = \frac{K^{CHS}_{cat,PCoA} CHS_t PCoA}{K^{PCoA}_{M,CHS} + PCoA}$$

---

##### CHI

$$v^{CHI}_{nar} = \frac{K^{CHI}_{cat,cha} CHS_t cha}{K^{cha}_{M,CHI} + cha}$$

---

### F3H

$$v^{F3H}_{DHK} = \frac{K^{F3H}_{cat,nar} F3H_t nar}{K^{nar}_{M,F3H} + nar}$$

---

### F3'H

$$v^{F3'H}_{DHQ} = \frac{K^{F3'H}_{cat,DHK} F3'H_t DHK}{K^{DHK}_{M,F3'H} + DHK}$$

---

**F3’5’H**

$$v_{DHM}^{F3'5'H} = \frac{K_{cat,DHQ}^{F3'5'H} F3'5'H_t DHQ}{K_{M,F3'5'H}^{DHQ} + DHQ}$$

---

**FLS**

$$v_{kam}^{FLS} = \frac{K_{cat,DHK}^{FLS} FLS_t DHK}{K_{M,FLS}^{DHK}(1 + \frac{DHQ}{K_{M,FLS}^{DHQ}} + \frac{DHM}{K_{M,FLS}^{DHM}}) + DHK}$$

$$v_{que}^{FLS} = \frac{K_{cat,DHQ}^{FLS} FNS_t DHQ}{K_{M,FLS}^{DHQ}(1 + \frac{DHK}{K_{M,FLS}^{DHK}} + \frac{DHM}{K_{M,FLS}^{DHM}}) + DHQ}$$

$$v_{myr}^{FLS} = \frac{K_{cat,DHM}^{FLS} FNS_t DHM}{K_{M,FLS}^{DHM}(1 + \frac{DHQ}{K_{M,FLS}^{DHQ}} + \frac{DHK}{K_{M,FLS}^{DHK}}) + DHM}$$

---

**DFR**

$$v_{LCP}^{DFR} = \frac{K_{cat,DHK}^{DFR} DFR_t DHK}{K_{M,DFR}^{DHK}(1 + \frac{DHQ}{K_{M,DFR}^{DHQ}} + \frac{DHM}{K_{M,DFR}^{DHM}}) + DHK}$$

$$v_{LCC}^{DFR} = \frac{K_{cat,DHQ}^{DFR} DFR_t DHQ}{K_{M,DFR}^{DHQ}(1 + \frac{DHK}{K_{M,DFR}^{DHK}} + \frac{DHM}{K_{M,DFR}^{DHM}}) + DHQ}$$

$$v_{LCD}^{DFR} = \frac{K_{cat,DHM}^{DFR} DFR_t DHM}{K_{M,DFR}^{DHM}(1 + \frac{DHQ}{K_{M,DFR}^{DHQ}} + \frac{DHK}{K_{M,DFR}^{DHK}}) + DHM}$$

---

**ANS**

$$v_{pel}^{ANS} = \frac{K_{cat,LCP}^{ANS} ANS_t LCP}{K_{M,ANS}^{LCP}(1 + \frac{LCC}{K_{M,ANS}^{LCC}} + \frac{LCD}{K_{M,ANS}^{LCD}}) + LCP}$$

$$v_{cya}^{ANS} = \frac{K_{cat,LCC}^{ANS} ANS_t LCC}{K_{M,ANS}^{LCC}(1 + \frac{LCP}{K_{M,ANS}^{LCP}} + \frac{LCD}{K_{M,ANS}^{LCD}}) + LCC}$$

$$v_{del}^{ANS} = \frac{K_{cat,LCD}^{ANS} ANS_t LCD}{K_{M,ANS}^{LCD}(1 + \frac{LCC}{K_{M,ANS}^{LCC}} + \frac{LCP}{K_{M,ANS}^{LCP}}) + LCD}$$

---

### Why do we see both regulatory and kinetic mutations in DFR contributing to shifts in the type of pigment produced?

Changes in DFR concentration can result in differential changes in flux down the anthocyanin branches. This arises for two reasons. The first reason is that pathway model is inherently biased toward the production of pelargoniding, followed by cyanidin, and then delphinidin. The reason for this bias is the ordering of branches. In the naive starting state, wherein all kinetic parameters are equivalent for all enzymes in the model, the pelargonidin branch receives 1/3 of the flux through the pathway up until that first branch point. The other 1/3 is then subsequently split among consecutive branches and so on due to the conservation of total flux through the pathway (Fig. S1). Thus, a larger share of the total flux contributes to the pelargonidin production than to production of the other anthocyanins. Increasing DFR concentration accentuates this bias, leading to a saturing curve (Fig. S4).

The second reason that DFR concentration changes contribute to branch-specificity is that when the activity of the DFR enzyme varies between precursors it can accentuate or overcome the bias of the pathway at any given DFR concentration as generalized in the previous section (Fig. S4). For example, a drastic reduction in the activity of DFR on the DHQ precursor (that leads to cyanidin production) will result increased pathway flux down the other branches. A simultaneous decrease in DFR concentration would lower the flux down the cyanidin branch to a greater extent than the other branches due to the already reduced activity of the reaction that leads to cyanidin production. This general effect of different parameter combinations on enzyme activity is shown in Fig. S5. The shape of these effect curves varies depending on the parameter that is altered to achieve different specificity. $K_{cat}$ and $E_t$ both result in linear increases in reaction rate, while $K_M$ has a non-linear, saturating effect. This is one underlying cause of extensive epistasis in the branched pathway model. The state of the kinetics of an enzyme determines the effects of changes to the concentration or to other kinetic parameters.

### Supplementary methods

#### Analysis of the simulated data

All simulations and analyses were performed in Python 3.6.7 using the following dependencies: numpy version 1.18.1, tellurium version 2.1.5, pandas version 0.25.3, matplotlib version 3.1.2, and seaborn version 0.9.0.

##### Calculation of overall parameter fixation frequencies, regulatory vs. biochemical parameter fixation frequencies, and normalized directional shifts

To identify hotspot loci in our simulations, we aggregated all the simulated data for each set of 10,000 unique trajectories for the two sequential phenotypic transitions. We then calculated the total number of fixation events involving each enzyme by summing over the constituent parameters. We also calculated an additional metric of fixation frequency, normalized to the number of unique parameters for each enzyme. The probability of randomly hitting an enzyme is proportional to the number of unique parameters for that enzyme, thus a per-parameter fixation rate ensures that comparisons between enzymes are made on the same shared scale. These values were further divided by the parameter type. We considered those that change enzyme function ($K_{cat}$ or $K_m$) as biochemical mutations and those that change enzyme amount ($E_t$) as regulatory.

To examine the dependence of the mutational patterns at hotspots on the phenotypic transition under selection, we also calculated a "normalized directional mutation size" ($\Delta$) for all mutations handled by the evolutionary algorithm. This value is calculated by taking the ratio: $(mutant\, value - previous\, value)/(previous\, value)$, yielding a fractional shift with a positive sign for mutations that increase the parameter value and a negative sign for mutations that decrease the parameter value. $\Delta$ is used to calculate additional informative values in subsequent analyses.

##### Analysis of selection coefficient curves, sensitivities, and fixation timing

To examine the distribution of selection coefficients for fixed mutations over the course of trajectories, we aggregated the selection coefficients for all fixed mutations for each set of 10,000 unique trajectories and stratified them by the trajectory step at which each mutation was fixed. To examine the selection effect per unit of parameter shift for each model parameter we logged the selection coefficient (s) and normalized directional mutation size ($\Delta$) for every individual mutation handled by the evolutionary algorithm (that did not drive the total steady state concentration outside the tolerated limit), not only the values for fixed parameters. We then calculated a "sensitivity" value ($|s/\Delta|$) for all of the pathway parameters across all mutations. This value captures how large of a shift in fitness is achieved by each mutation of a given size. These normalized sensitivities provided a metric to compare the differential effects of mutations at different pathway parameters. Finally, we aggregated the data for the timing of fixations at all pathway parameters over each set of 10,000 trajectories, expressing these data as the trajectory step at which each fixation event occurred. This allowed us to examine different stepwise distributions of fixation events across the set of pathway parameters.

##### Analysis of shifts in pathway dynamics and control structure

To examine the structure of control over pathway dynamics by the component enzymes in our model, we used the metabolic control analysis (MCA) tools available in the Tellurium python environment for metabolic simulations (Choi et al., 2018). MCA provides a set of mathematical tools for studying how individual reactions control the flux of material through integrated metabolic networks (Cornish-Bowden, 1995; Kacser et al., 1995). For each starting-state and end-state, we calculated the concentration control coefficients for each pathway reaction over each chemical species concentration at steady state. These control coefficients, the elements of the Jacobian matrix for the pathway dynamics, represent the partial derivatives of each steady state concentration with respect to the activity of each reaction (Cornish-Bowden, 1995; Kacser et al., 1995). They capture the ability of the individual reactions to control the pathway output. Changes in the space of control coefficients is directly tied to the shifts in pathway flux that result in phenotypic transitions. Therefore, examining this pathway "control structure" provides a detailed way of examining how the pathway flux has been tuned during simulated evolution. For all analyses of control coefficients we excluded the effects of the fixed $K_{sink}$ rate constant for the boundary transport process of pathway products.

### Availability of data and materials

The python library *enzo*, written to perform evolutionary simulations of the pathway model, is available on github (https://github.com/lcwheeler/enzo). The scripts used to run the simulations and Jupyter notebooks and scripts for the subsequent analyses, and the raw simulated datasets for the main simulations ($naive-to-delphinidin-simulated-dataset.p$, $delphinidin-to-cyanidin-simulated-dataset.p$, and $cyanidin-to-pelargonidin-simulated-dataset.p$) are available in a OSF online repository (https://osf.io/kxr23/). The simulated dataset files ($\sim 5GB$) are presented in serialized (pickle) format (the entire *enzo* PathwaySet simulation objects are pickled in this manner) and have been zipped. To use these pickled (".p") dataset files they must first be unzipped.

### Supplemental figures

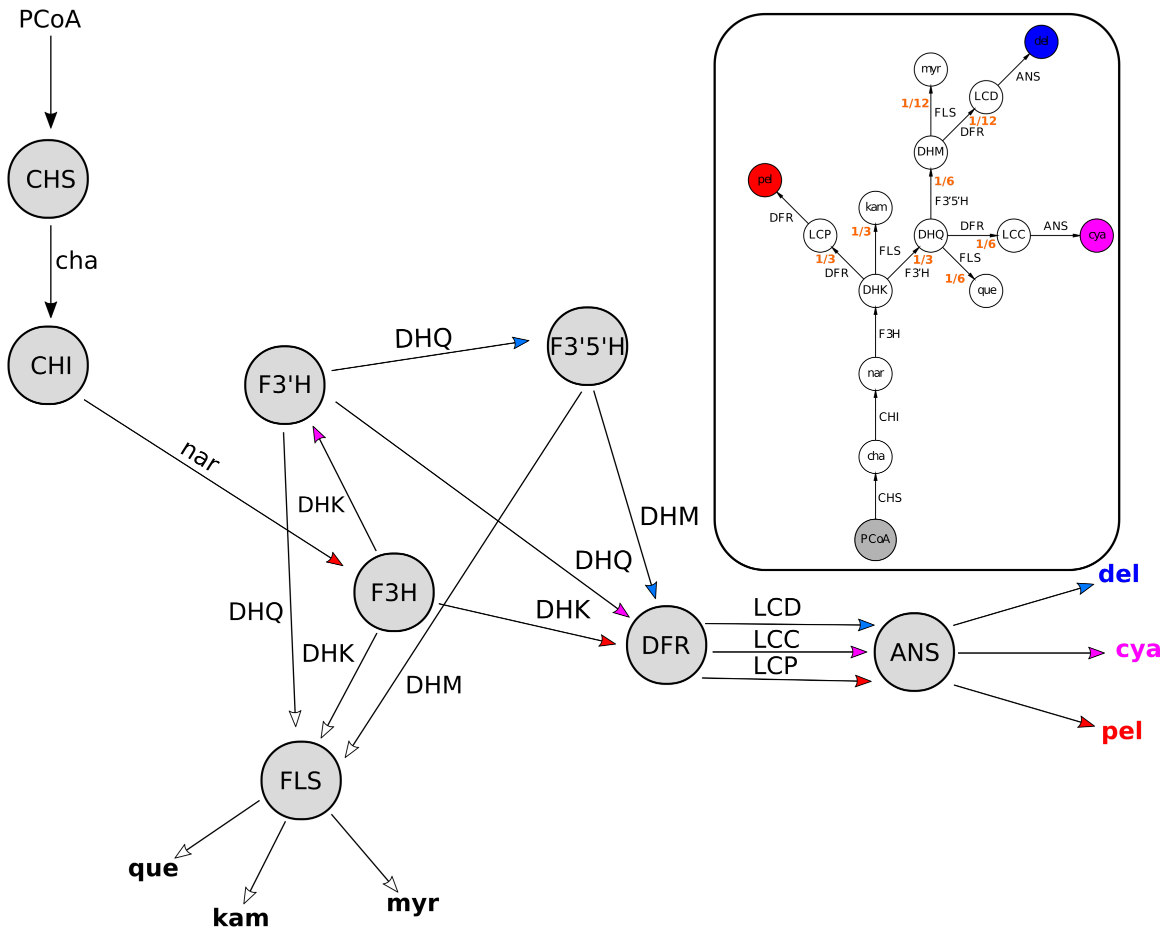


**Fig S1. Alternate diagrams of the anthocyanin pathway.** Enzyme-centric pathway diagram is shown. In this representation circles are enzymes and arrows are substrates/reactions (labeled). This view helps to highlight the competition between substrates for a single pool of enzyme. The inset shows the substrate-centric diagram from Fig. 1 in the main text, with the partitioning of flux down the pathway branches that occurs in the naive model labeled in orange. Floating species abbreviations are: PCoA (P-Coumaroyl-CoA), cha (chalcone), nar (naringenin), DHK (dihydrokampferol), DHQ (dihydroquercetin), DHM (dihydromyricetin), que (quercetin), kam (kampferol), myr (myricetin), LCD (leucopelargonidin), LCC (leucocyanidin), LCD (leucodelphinidin), pel (pelargonidin), cya (cyanidin), del (delphinidin). Del, cya, and pel are the anthocyanidins, which are glycosylated to form the various anthocyanins. Kam, que, and myr are the flavonols. Enzyme abbreviations are: CHS (chalcone synthase), CHI (chalcone isomerase), F3H (flavanone-3-hydroxylase), F3'H (flavonol-3'hydroxylase), F3'5'H (flavonoid-3'5'hydroxylase), DFR (dihyroflavonol-4-reductase), FLS (flavonol synthase), ANS (anthocyanidin synthase).

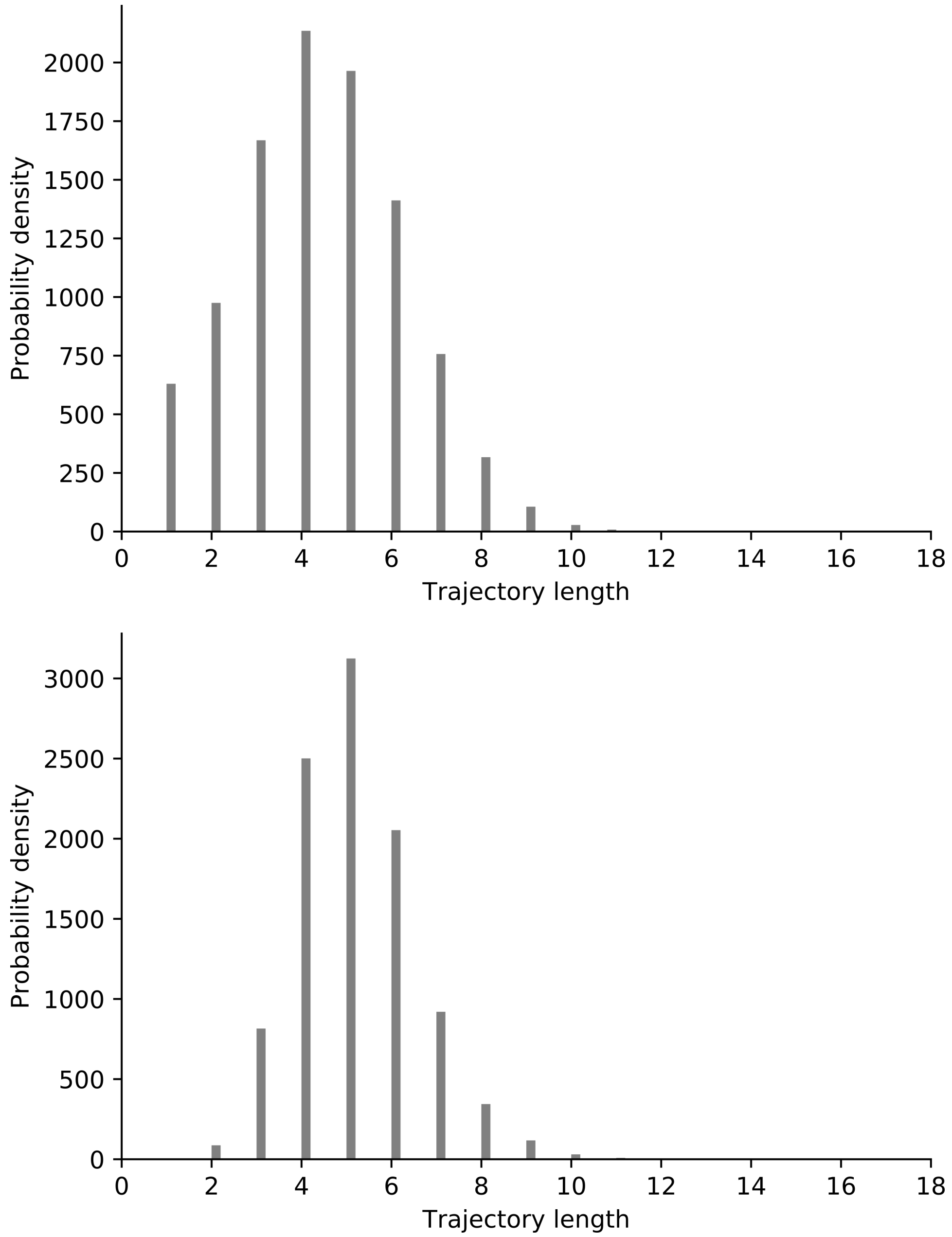


**Fig S2. Distributions of simulated trajectory lengths.** Histograms show the distributions of trajectory lengths in mutational steps for the delphinidin to cyanidin transition (top) and the cyanidin to pelargonidin transition (bottom).

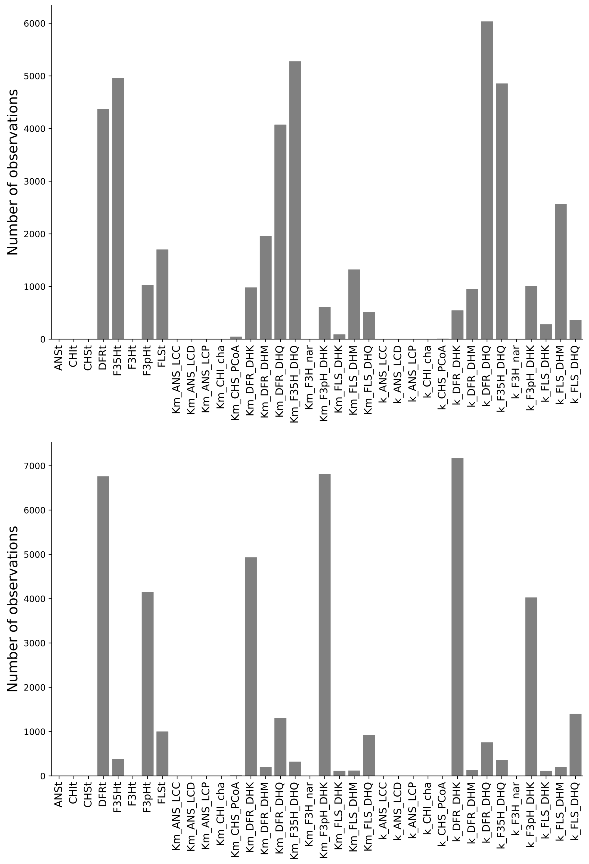


**Fig S3. Fixation events per evolvable model parameter.** Total fixed mutation counts for each evolvable model parameter for the delphinidin to cyanidin transition (top) and the cyanidin to pelargonidin transition (bottom).

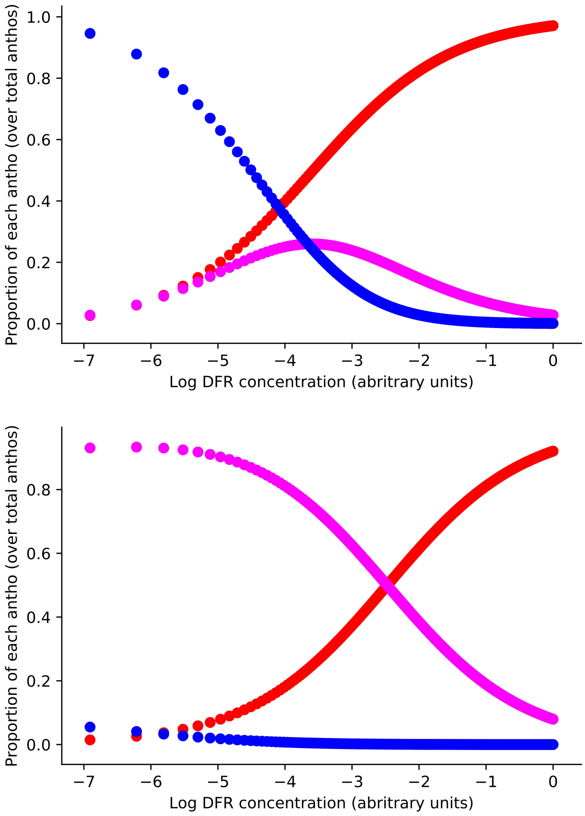


**Fig S4. Effects of DFR concentration changes on anthocyanin output.** Top: delphinidin starting state. Bottom: cyanidin starting state. Plots show the effect of increasing DFR concentration on the proportion of the total anthocyanin content at steady state comprised by each pigment, in both of the starting state models used for sequential phenotypic transitions. As DFR concentration becomes large the pathway produces primarily pelargonidin, even in the evolved states optimized for production of delphinidin or cyanidin. This effect is due to the early partitioning of a large portion of flux down the path leading to pelargonidin (see Fig. S1).

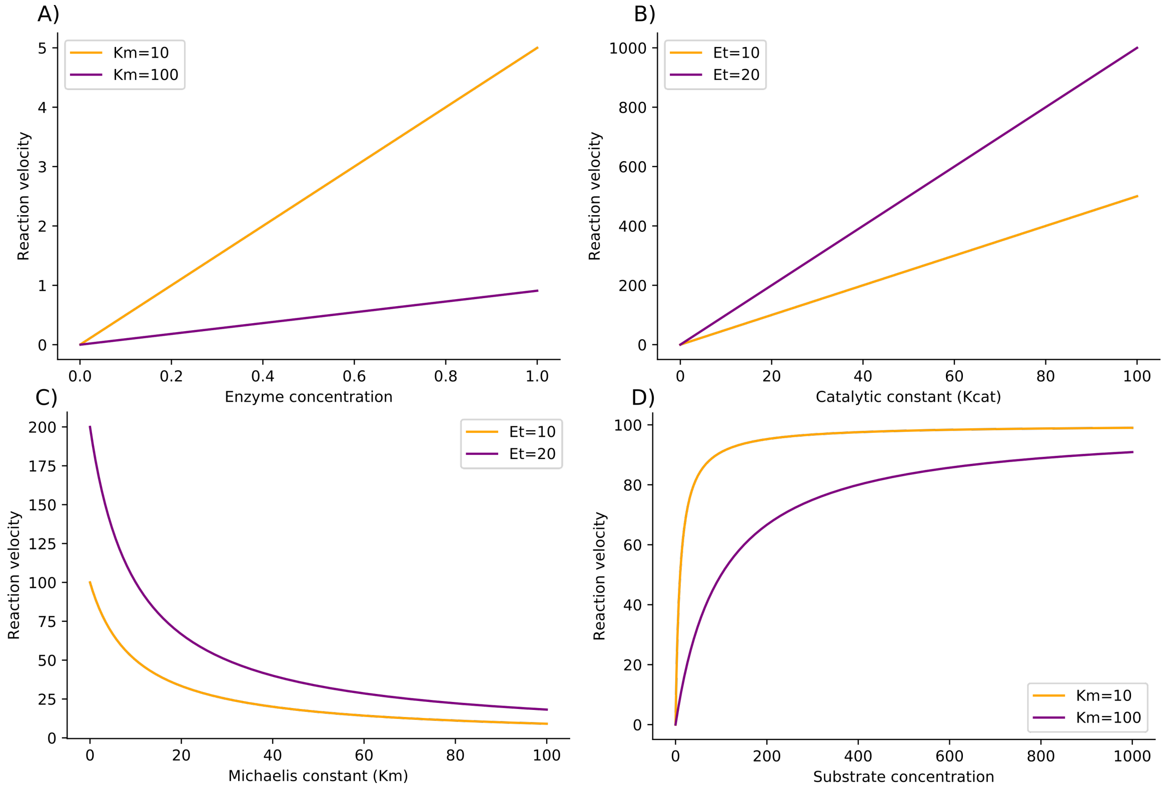


**Fig S5. General effects of kinetic differences between enzymes.** A simple one-enzyme, one-substrate system shown initialized with two different parameter values for catalytic constant ($K_{cat}$), Michaelis constant ($K_m$), and enzyme concentration ($E_t$). The value of enzyme parameters causes differential effects on reaction velocity ($v$) when introducing changes at another enzyme parameter or scanning across another pathway variable. A) Effect of varying enzyme concentration at two different values of $K_m$. B) Effect of varying $K_{cat}$ at two different values of $E_t$. C) Effect of varying $K_m$ at two different values of $E_t$. D) Effect of varying substrate concentration (S) at two different values of $K_m$.

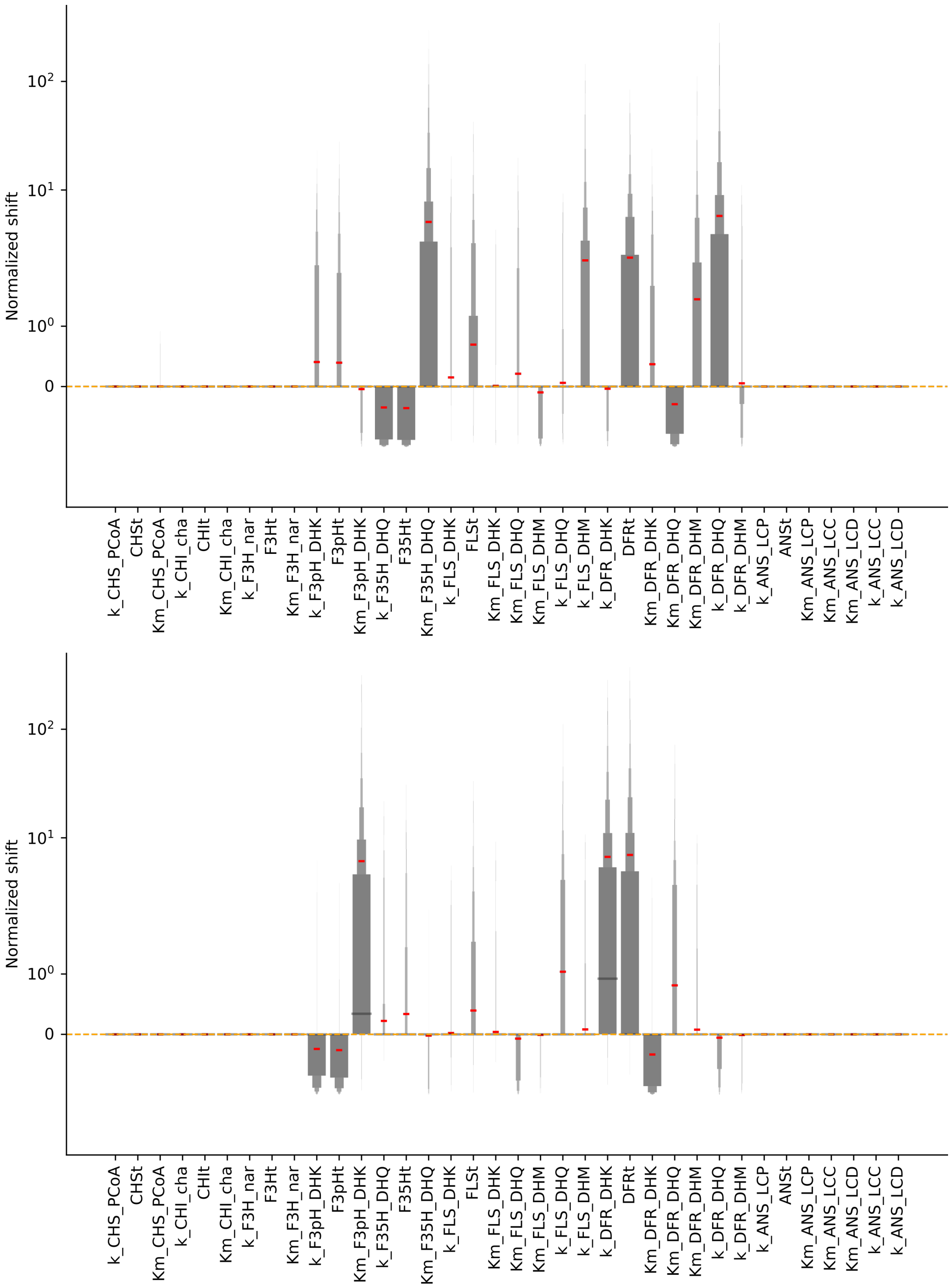


**Fig S6. End-state normalized directional shifts for all evolvable parameters.** Normalized directional shift distributions for all evolvable model parameters, shown as boxenplots (outliers not shown) for the delphinidin to cyanidin transition (top) and the cyanidin to pelargonidin transition (bottom).

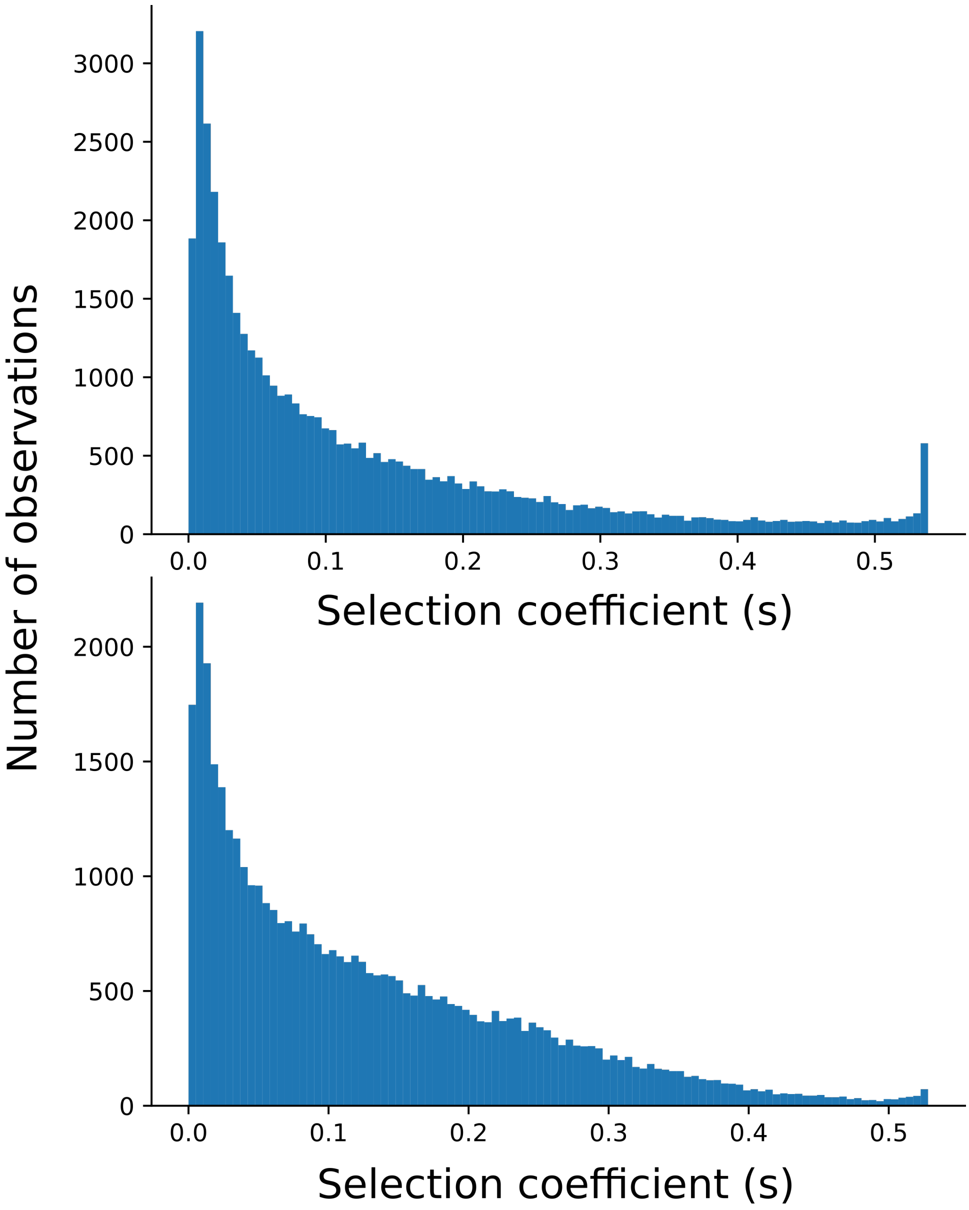


**Fig S7. Aggregated distributions of fixed selection coefficients.** Histograms show aggregated distributions of fixed selection coefficients for the delphinidin to cyanidin transition (top) and the cyanidin to pelargonidin transition (bottom).

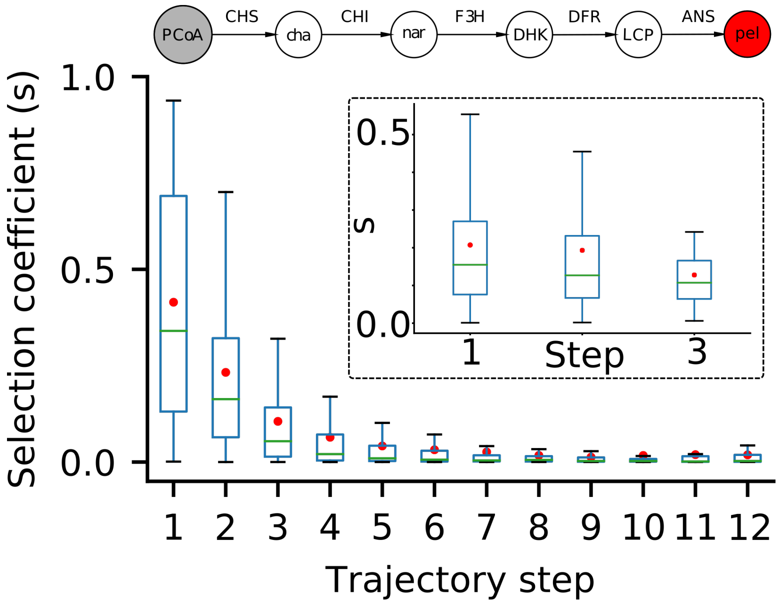


**Fig S8. A simple linear model exhibits monotonically-decreasing selection coefficients over trajectory steps.** Stepwise distributions of fixed selection coefficients for trajectories from 2,000 simulations of a linear pathway (shown at the top of figure). Main plot shows all trajectories, including the 43% that did not find the phenotypic optimum before the alloted number of iterations was reached. Inset shows distributions only for the 57% of trajectories that did reach the optimum (all of which were no more than 3 steps long). Distributions are shown as boxplots. Green lines are median, red dots are mean.

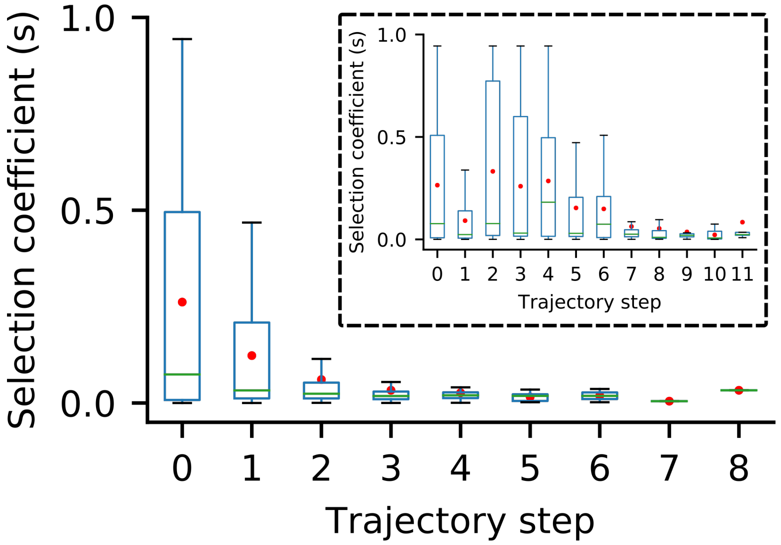


**Fig S9. Altering the fitness function has a strong affect on the stepwise distribution of selection coefficients.** Stepwise distributions of fixed selection coefficients for trajectories from 2,000 simulations of the naive pathway model under an alternative fitness function: stabilizing selection on ratio of pelargonidin to total steady state concentration of all floating species, with optimum at 3-fold increase in absolute pelargonidin concentration. Main plot shows distributions only for the 99.3% of trajectories that reached the phenotypic optimum. Inset shows all trajectories, including the 0.7% that did not reach the optimum before the alloted number of iterations was reached. These long trajectories strongly skew the distributions at each step. Distributions are shown as boxplots. Green lines are median, red dots are mean.

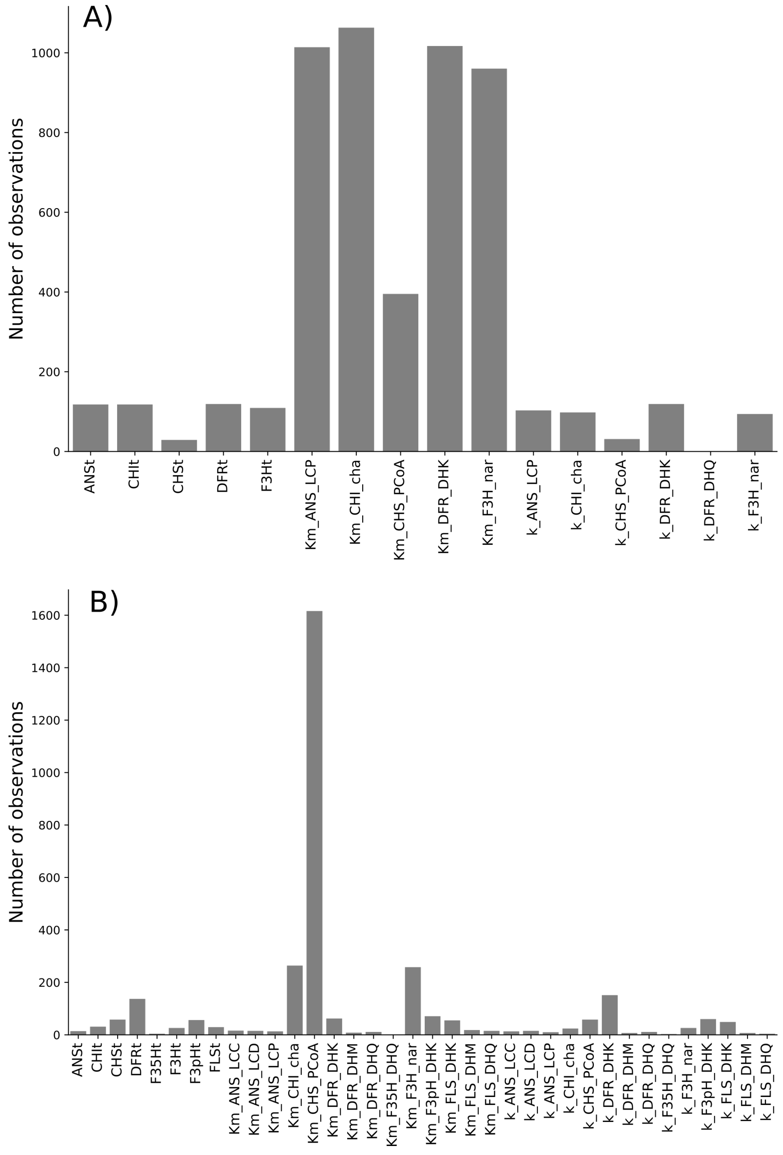


**Fig S10. Altering the fitness function and pathway topology results in different pathway hotspots.** Total fixation counts at each model parameter across all 2,000 control simulations for A) simplified lineary pathway (see Fig. S6) and B) naive anthocyanin pathway model. Fitness function is stabilizing selection on the ratio of pelargonidin to total steady state concentration of all floating species, with optimum at 3-fold increase in absolute pelargonidin concentration.

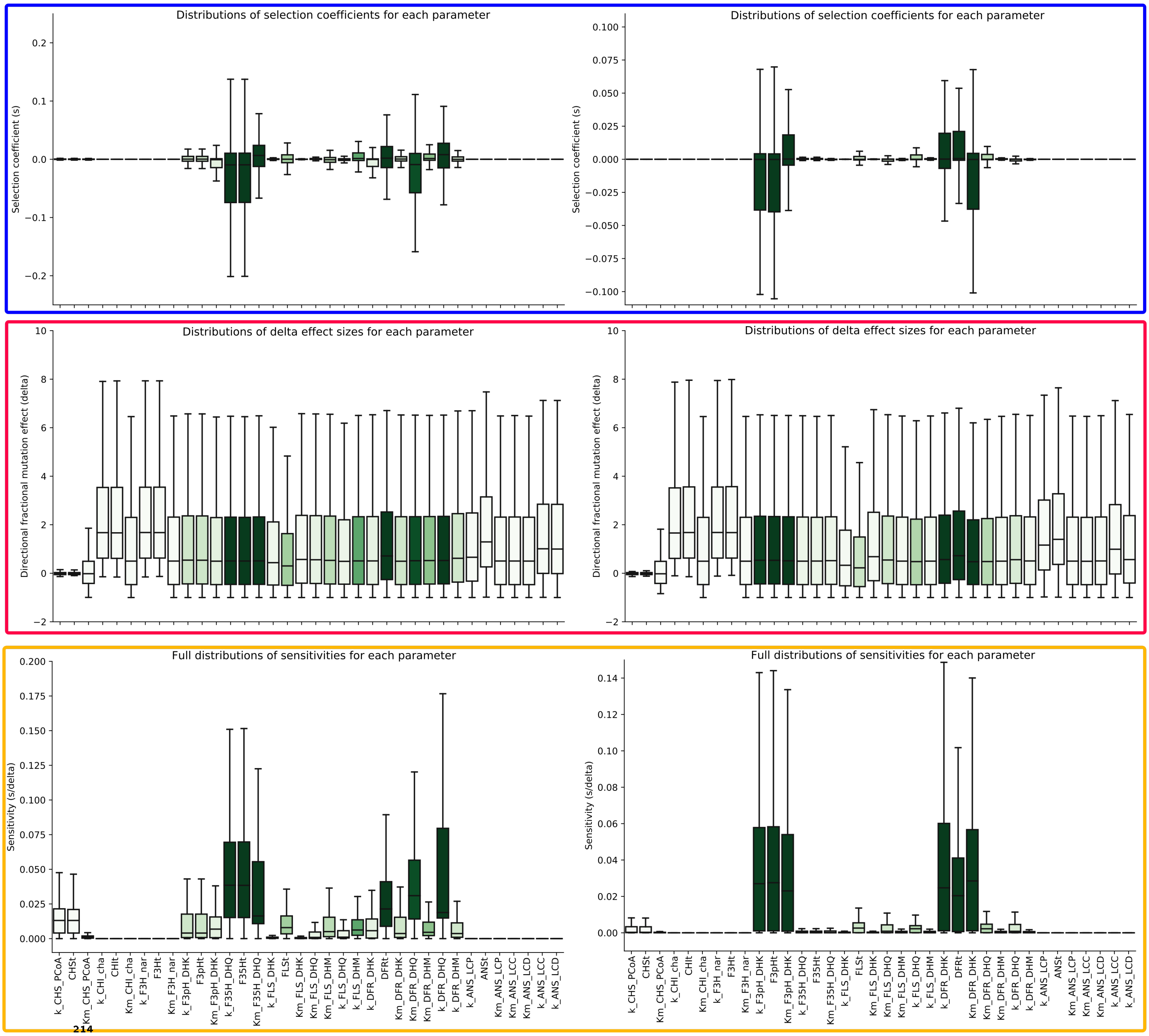


**Fig S11. Sensitivity values for each parameter calculated from selection coefficients and normalized directional mutation sizes.** Left column: 90% delphinidin to 90% cyanindin. Right column: 90% cyanidin to 90% pelargonidin. Distributions are depicted by boxplots, colored by the number of fixation events ranging from 0 (white) to approximately 8000 (dark green). Results shown for all parameter mutations handled by the algorithm (except for those that push the total steady state concentraton outside tolerance envelope). Blue box: Selection coefficients (s). Red box: normalized directional mutation size ($\Delta$): $(mutant\,value - previous\,value)/(previous\,value)$).

Orange box: sensitivity values calculated by dividing the s values in the blue box by the $\Delta$ values in the red box for every mutation, then taking the absolute value ($|s/\Delta|$).

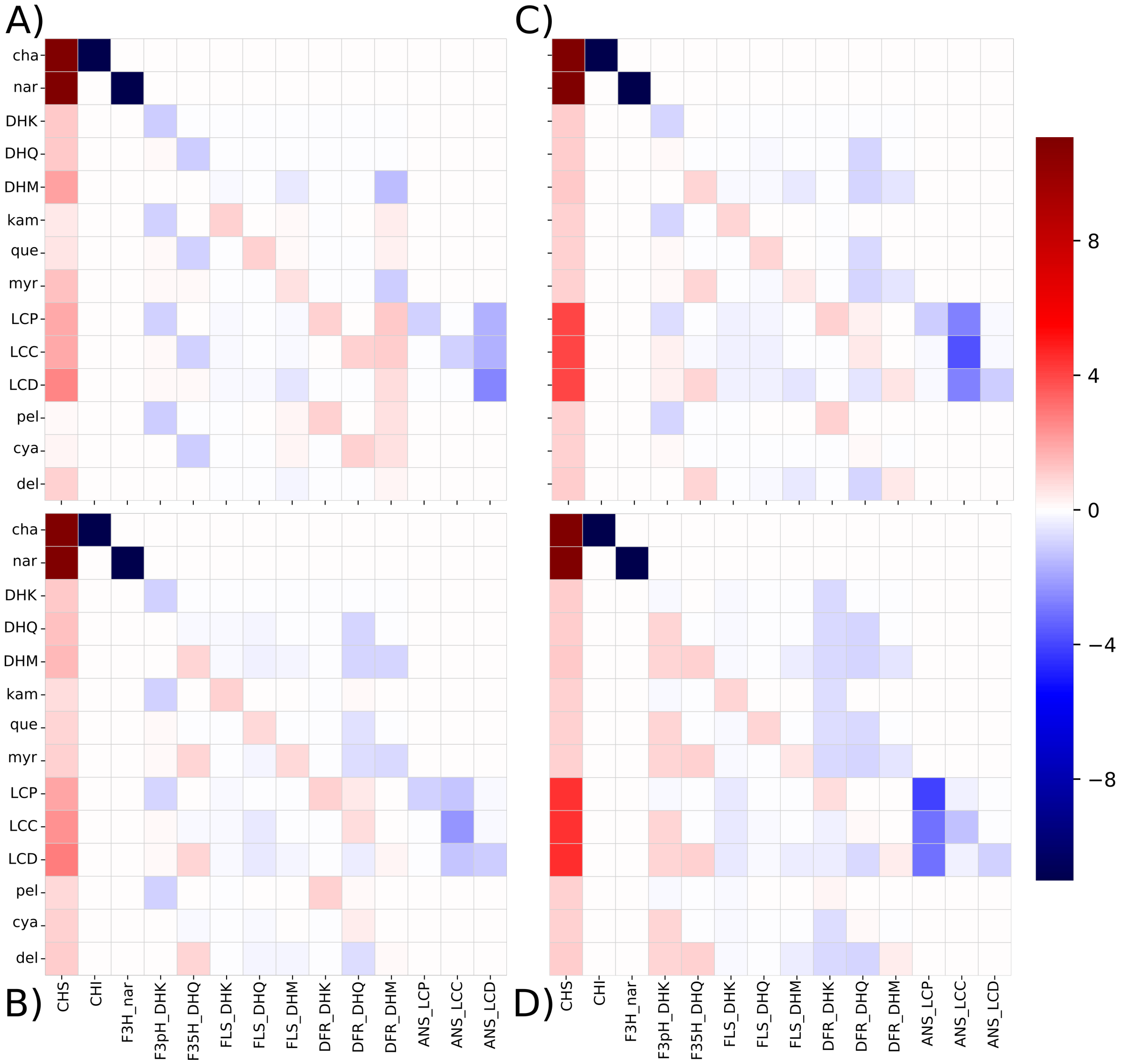


**Fig S12. Matrices of concentration control coefficients for the median evolved states.** Heatmaps depict the concentration control coefficients for each pathway reaction on the steady state concentration of each floating chemical species in the model. Left column shows transtion from delphinidin starting state (A) to cyanidin end state (B). Right column shows cyanidin starting state (C) to pelargonidin end state (D).

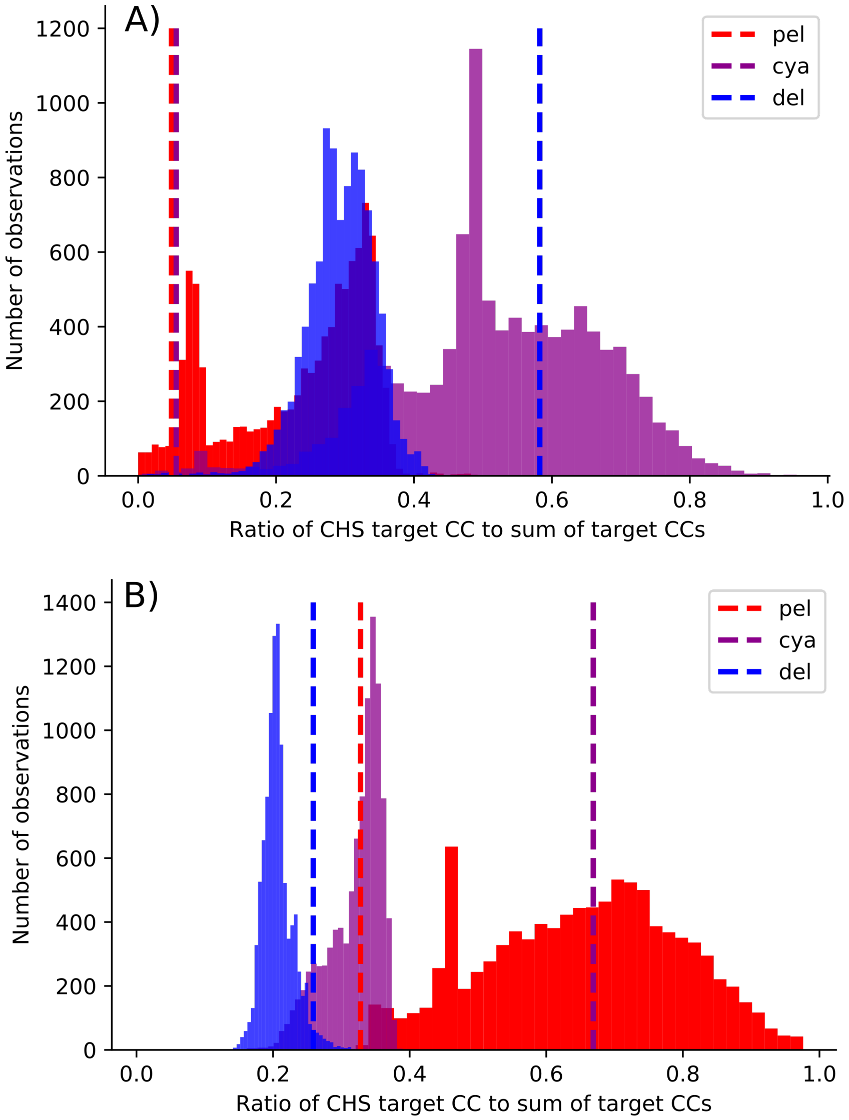


**Fig S13. Distributions of relative CHS control for evolutionary end-states.** Histograms show the absolute valye of the ratio of CHS control coefficient for each anthocyanin pigment to the sum of all control coefficients for the same pigment ($|CC^{CHS}_{target}/\sum CC^{all\,enzymes}_{target}|$) across evolutionary end-states from all trajectories for (A) delphinidin starting state to cyanidin end state and (B) cyanidin starting state to pelargonidin end state. Dashed vertical lines indicate the value of $|CC^{CHS}_{target}/\sum CC^{all\,enzymes}_{target}|$ in the starting state of the evolutionary simulations.

| Citation | Taxon | Color change | Pigment change | Enzyme Changes | Type |
|---|---|---|---|---|---|
| (Hoshino et al., 2003) | *Ipomoea nil, I. tricolor,I. purpurea* | blue/red polymorphism | cya to pel | F3'H (coding mutations) | N |
| (Zufall and Rausher, 2004) | *Ipomoea quamoclit* | blue to red | cya to pel | F3'H (downregulation), DFR (specificity) | N |
| (Streisfeld and Rausher, 2009) | *Ipomoea udeana, I. quamoclit, I. horsfalliae* | blue to red | cya to pel | F3'H (downregulation) | N |
| (Marais and Rausher, 2010) | *Ipomoea purpurea, I, quamoclit, I. coccinea, I. ternifolia* | blue to red/pink | cya to pel | F3'H (downregulation) | N |
| (Hopkins and Rausher, 2011) | *Phlox drummondii* | blue to red | del to cya | F3'5'H (downregulation) | N |
| (Mizuta et al., 2010) | *'Oomurasaki' azalea* | purple/red polymorphism | del to cya | F3'5'H (downregulation) | N |
| (Ishiguro et al., 2012) | *Antirrhinum kelloggii* | blue to red | del to cya | F3'H (downregulation) | N |
| (Nakatsuka et al., 2006) | *Gentiana scabra* | blue to pink | del to cya | F3'5'H (deactivation) | N |
| (Matsubara et al., 2005) | *Petunia hybrida* | blue to red/pink | del to cya | F3'5'H (deactivation) | N |
| (Mizuta et al., 2014) | *Rhododendron kiusianum, Rhododendron kaempferi* | red/purple polymorphism | del to cya | F3'5'H (downregulation) | N |
| (Moreau et al., 2012) | *Pisum sativum* | purple to pink | del to cya | F3'5'H (deactivation) | N |
| (Wessinger and Rausher, 2014) | *Penstemon barbatus* | blue to red | del to pel | F3'5'H (downregulation) | N |
| (Wessinger and Rausher, 2015) | *Penstemon labrosus, P. subulatus, P. rostrifloris, P. baccharifolius, P. alamosensis, P. miniatus, P. superbus, P. murrayanus, P. centranthifolius, P. utahensis, P. pinifolius, P. eatonii* | blue to red | del to pel | F3'5'H (degeneration/deactivation) | N |
| (Smith and Rausher, 2011) | *Iochroma gesnerioides* | blue to red | del to pel | F3'H (downregulation), F3'5'H (deletion),DFR (specificity) | N |
| (Freyre et al., 2015) | *Ruellia simplex* | pink/purple polymorphism | pel to del | F3'H (upregulation), F3'5'H (upregulation) | N |
| (Sato et al., 2011) | *Saintpaulia* sp. | purple/pink/blue polymorphism | pel/cya/del variation | F3'5'H (regulatory mutations) | E |
| (Nakatsuka et al., 2007) | *Nicotiana tabacum* | pink to red | cya to pel | FLS (downregulation), F3'H (downregulation), DFR (specificity/transgene expression) | E |
| (Meyer et al., 1987) | *Petunia hybrida* | pink to red | cyanidin to pelargonidin | DFR (specificity/transgene expression) | E |
| (Nakamura et al., 2010) | *Torenia hybrida* | blue to pink | del to pel | F3'5'H (downregulation), F3'H (downregulation), DFR (specificity/transgene expression) | E |
| (Seitz et al., 2007) | *Osteospermum hybrida* | blue to red | del to pel | F3'5'H (downregulation) | E |
| (Noda et al., 2013) | *Chrysanthemum morifolium* | red to violet | pel/cya to del | F3'5'H (upregulated transgene) | E |
| (Fukui et al., 2003) | *Dianthus caryophyllus* | pink to blue | pel to del | F3'5'H (upregulated transgene) | E |
| (Nakamura et al., 2015) | *Rosa hybrida* | pink to purple | pel to cya/del | F3'5'H (upregulated transgene) | E |
| (Katsumoto et al., 2007) | *Rosa hybrida* | red to blue | pel/cya to del | F3'5'H (upregulated transgene),DFR (specificity/transgene expression) | E |
| (Nakatsuka et al., 2008) | *Gentiana triflora, Gentiana scabra* | blue to magenta | del to cya | F3'5'H (downregulation) | E |
| (Ueyama et al., 2002) | *Torenia hybrida* | blue to magenta | del to cya | F3'H (downregulation), FNS (downregulation) | E |
| (Tsuda et al., 2004) | *Petunia hybrida* | purple to pink | del to cya | F3'H (downregulation) | E |
| (Boase et al., 2010) | *Cyclamen persicum* | purple to red/pink | del to cya | F3'5'H (downregulation) | E |
| (Nielsen et al., 2002) | *Eustoma grandiflorum* | blue to purple | del to cya | FLS (upregulated transgene) | E |
| (Shimada et al., 2001) | *Petunia hybrida* | pink to magenta | cya to del | F3'5'H (upregulated transgene) | E |
| (Okinaka et al., 2003) | *Nicotiana tabacum* | pink to purple | cya to del | F3'5'H (upregulated transgene) | E |
| (Brugliera et al., 2013) | *Chrysanthemum× morifolium Ramat.* | pink to purple-blue | cya to del | F3'5'H (upregulated transgene) | E |

**Table S1. Empirical studies of the genetic basis for differences in floral anthocyanin type in natural and engineered systems.** This table is reproduced from the one published previously (Wheeler and Smith, 2019). Natural (N) examples encompass changes that have occurred in the wild along with spontaneous changes in cultivated varieties. Engineered (E) changes are those that have been intentially introduced using biotechnology (i.e. transgenics, gene silencing, etc.). Pigments are abbreviated: pelargonidin (pel), cyanidin (cya), and delphinidin (del) as in figure S1.

### References


Boase, M. R., Lewis, D. H., Davies, K. M., Marshall, G. B., Patel, D., Schwinn, K. E., and Deroles, S. C. Isolation and antisense suppression of flavonoid 3’, 5’-hydroxylasemodifies flower pigments and colour in cyclamen. *BMC Plant Biology*, 10(1):107, June 2010. ISSN 1471-2229.

Brugliera, F., Tao, G.-Q., Tems, U., Kalc, G., Mouradova, E., Price, K., Stevenson, K., Nakamura, N., Stacey, I., Katsumoto, Y., Tanaka, Y., and Mason, J. G. Violet/blue chrysanthemums—metabolic engineering of the anthocyanin biosynthetic pathway results in novel petal colors. *Plant Cell Physiol*, 54(10):1696–1710, Oct. 2013. ISSN 0032-0781.

Choi, K., Medley, J. K., König, M., Stocking, K., Smith, L., Gu, S., and Sauro, H. M. Tellurium: An extensible python-based modeling environment for systems and synthetic biology. *Biosystems*, 171:74–79, 2018. ISSN 0303-2647. doi: https://doi.org/10.1016/j.biosystems.2018.07.006.

Cornish-Bowden, A. Metabolic control analysis in theory and practice. In Bittar, E. E., editor, *Advances in Molecular and Cell Biology*, volume 11 of *Enzymology in Vivo*, pages 21–64. Elsevier, Jan. 1995.

Freyre, R., Uzdevenes, C., Gu, L., and Quesenberry, K. H. Genetics and anthocyanin analysis of flower color in mexican petunia. *J. Am. Soc. Hortic. Sci.*, 140(1):45–49, Jan. 2015. ISSN 2327-9788, 0003-1062.

Fukui, Y., Tanaka, Y., Kusumi, T., Iwashita, T., and Nomoto, K. A rationale for the shift in colour towards blue in transgenic carnation flowers expressing the flavonoid 3′,5′-hydroxylase gene. *Phytochemistry*, 63(1):15–23, May 2003. ISSN 0031-9422.

Hopkins, R. and Rausher, M. D. Identification of two genes causing reinforcement in the texas wildflower *Phlox drummondii*. *Nature*, 469(7330):411–414, Jan. 2011. ISSN 1476-4687.

Hoshino, A., Morita, Y., Choi, J.-D., Saito, N., Toki, K., Tanaka, Y., and Iida, S. Spontaneous mutations of the flavonoid 3′-hydroxylase gene conferring reddish flowers in the three morning glory species. *Plant Cell Physiol*, 44(10):990–1001, Oct. 2003. ISSN 0032-0781.

Ishiguro, K., Taniguchi, M., and Tanaka, Y. Functional analysis of antirrhinum kelloggii flavonoid 3′-hydroxylase and flavonoid 3′,5′-hydroxylase genes; critical role in flower color and evolution in the genus antirrhinum. *J Plant Res*, 125(3):451–456, May 2012. ISSN 1618-0860.

Kacser, H., Burns, J. A., Kacser, H., and Fell, D. A. The control of flux. *Biochem. Soc. Trans.*, 23 (2):341–366, May 1995. ISSN 0300-5127, 1470-8752.

Katsumoto, Y., Fukuchi-Mizutani, M., Fukui, Y., Brugliera, F., Holton, T. A., Karan, M., Nakamura, N., Yonekura-Sakakibara, K., Togami, J., Pigeaire, A., Tao, G.-Q., Nehra, N. S., Lu, C.-Y., Dyson, B. K., Tsuda, S., Ashikari, T., Kusumi, T., Mason, J. G., and Tanaka, Y. Engineering of the rose flavonoid biosynthetic pathway successfully generated blue-hued flowers accumulating delphinidin. *Plant Cell Physiol*, 48(11):1589–1600, Nov. 2007. ISSN 0032-0781.

Marais, D. L. D. and Rausher, M. D. Parallel evolution at multiple levels in the origin of hummingbird pollinated flowers in ipomoea. *Evolution*, 64(7):2044–2054, 2010. ISSN 1558-5646.

Matsubara, K., Kodama, H., Kokubun, H., Watanabe, H., and Ando, T. Two novel transposable elements in a cytochrome p450 gene govern anthocyanin biosynthesis of commercial petunias. *Gene*, 358:121–126, Sept. 2005. ISSN 0378-1119.

Meyer, P., Heidmann, I., Forkmann, G., and Saedler, H. A new petunia flower colour generated by transformation of a mutant with a maize gene. *Nature*, 330(6149):677, Dec. 1987. ISSN 1476-4687.

Mizuta, D., Nakatsuka, A., Miyajima, I., Ban, T., and Kobayashi, N. Pigment composition patterns and expression analysis of flavonoid biosynthesis genes in the petals of evergreen azalea 'oomurasaki' and its red flower sport. *Plant Breed.*, 129(5):558–562, 2010. ISSN 1439-0523.

Mizuta, D., Nakatsuka, A., Ban, T., Miyajima, I., and Kobayashi, N. Pigment composition patterns and expression of anthocyanin biosynthesis genes in rhododendron kiusianum, r. kaempferi, and their natural hybrids on kirishima mountain mass, japan. *J. Japan. Soc. Hort. Sci.*, pages CH–087, Mar. 2014. ISSN 1882-3351, 1882-336X.

Moreau, C., Ambrose, M. J., Turner, L., Hill, L., Ellis, T. H. N., and Hofer, J. M. I. The b gene of pea encodes a defective flavonoid 3′,5′-hydroxylase, and confers pink flower color. *Plant Physiol.*, 159(2):759–768, June 2012. ISSN 0032-0889, 1532-2548.

Nakamura, N., Fukuchi-Mizutani, M., Fukui, Y., Ishiguro, K., Suzuki, K., Suzuki, H., Okazaki, K., Shibata, D., and Tanaka, Y. Generation of pink flower varieties from blue torenia hybrida by redirecting the flavonoid biosynthetic pathway from delphinidin to pelargonidin. *Plant Biotechnol.*, 27(5):375–383, 2010. ISSN 1347-6114, 1342-4580.

Nakamura, N., Katsumoto, Y., Brugliera, F., Demelis, L., Nakajima, D., Suzuki, H., and Tanaka, Y. Flower color modification in *Rosa hybrida* by expressing the *S*-adenosylmethionine: anthocyanin 3′,5′-*O*-methyltransferase gene from *Torenia hybrida*. *Plant Biotechnol.*, 32(2):109–117, 2015. ISSN 1342-4580, 1347-6114.

Nakatsuka, T., Nishihara, M., Mishiba, K., Hirano, H., and Yamamura, S. Two different transposable elements inserted in flavonoid 3′,5′-hydroxylase gene contribute to pink flower coloration in gentiana scabra. *Mol Genet Genomics*, 275(3):231–241, Mar. 2006. ISSN 1617-4623.

Nakatsuka, T., Abe, Y., Kakizaki, Y., Yamamura, S., and Nishihara, M. Production of red-flowered plants by genetic engineering of multiple flavonoid biosynthetic genes. *Plant Cell Rep*, 26(11): 1951–1959, Nov. 2007. ISSN 1432-203X.

Nakatsuka, T., Mishiba, K.-i., Abe, Y., Kubota, A., Kakizaki, Y., Yamamura, S., and Nishihara, M. Flower color modification of gentian plants by rnai-mediated gene silencing. *Plant Biotechnol.*, 25 (1):61–68, 2008. ISSN 1347-6114, 1342-4580.

Nielsen, K., Deroles, S. C., Markham, K. R., Bradley, M. J., Podivinsky, E., and Manson, D. Antisense flavonol synthase alters copigmentation and flower color in lisianthus. *Molecular Breeding*, 9(4):217–229, Dec 2002. ISSN 1572-9788. doi: 10.1023/A:1020320809654.

Noda, N., Aida, R., Kishimoto, S., Ishiguro, K., Fukuchi-Mizutani, M., Tanaka, Y., and Ohmiya, A. Genetic engineering of novel bluer-colored chrysanthemums produced by accumulation of delphinidin-based anthocyanins. *Plant Cell Physiol*, 54(10):1684–1695, Oct. 2013. ISSN 0032-0781.

Okinaka, Y., Shimada, Y., Nakano-Shimada, R., Ohbayashi, M., Kiyokawa, S., and Kikuchi, Y. Selective accumulation of delphinidin derivatives in tobacco using a putative flavonoid 3′,5′-hydroxylase cdna from campanula medium. *Biosci. Biotechnol. Biochem.*, 67(1):161–165, 2003. ISSN 0916-8451, 1347-6947.

Sato, M., Kawabe, T., Hosokawa, M., Tatsuzawa, F., and Doi, M. Tissue culture-induced flower-color changes in saintpaulia caused by excision of the transposon inserted in the flavonoid 3′, 5′ hydroxylase (f3′5′h) promoter. *Plant Cell Rep*, 30(5):929–939, May 2011. ISSN 1432-203X.

Seitz, C., Vitten, M., Steinbach, P., Hartl, S., Hirsche, J., Rathje, W., Treutter, D., and Forkmann, G. Redirection of anthocyanin synthesis in osteospermum hybrida by a two-enzyme manipulation strategy. *Phytochemistry*, 68(6):824–833, Mar. 2007. ISSN 0031-9422.

Shimada, Y., Ohbayashi, M., Nakano-Shimada, R., Okinaka, Y., Kiyokawa, S., and Kikuchi, Y. Genetic engineering of the anthocyanin biosynthetic pathway with flavonoid-3′,5′-hydroxylase: specific switching of the pathway in petunia. *Plant Cell Rep*, 20(5):456–462, July 2001. ISSN 1432-203X.

Smith, S. D. and Rausher, M. D. Gene loss and parallel evolution contribute to species difference in flower color. *Mol. Biol. Evol.*, 28(10):2799–2810, Oct. 2011. ISSN 0737-4038.

Streisfeld, M. A. and Rausher, M. D. Genetic changes contributing to the parallel evolution of red floral pigmentation among ipomoea species. *New Phytol.*, 183(3):751–763, 2009. ISSN 1469-8137.

Tsuda, S., Fukui, Y., Nakamura, N., Katsumoto, Y., Yonekura-Sakakibara, K., Fukuchi-Mizutani, M., Ohira, K., Ueyama, Y., Ohkawa, H., Holton, T. A., Kusumi, T., and Tanaka, Y. Flower color modification of petunia hybrida commercial varieties by metabolic engineering. *Plant Biotechnol.*, 21(5):377–386, 2004. ISSN 1347-6114, 1342-4580.

Ueyama, Y., Suzuki, K.-i., Fukuchi-Mizutani, M., Fukui, Y., Miyazaki, K., Ohkawa, H., Kusumi, T., and Tanaka, Y. Molecular and biochemical characterization of torenia flavonoid 3′-hydroxylase and flavone synthase ii and modification of flower color by modulating the expression of these genes. *Plant Science*, 163(2):253–263, Aug. 2002. ISSN 0168-9452.

Wessinger, C. A. and Rausher, M. D. Predictability and irreversibility of genetic changes associated with flower color evolution in penstemon barbatus. *Evolution*, 68(4):1058–1070, 2014. ISSN 1558-5646.

Wessinger, C. A. and Rausher, M. D. Ecological transition predictably associated with gene degeneration. *Mol. Biol. Evol.*, 32(2):347–354, Feb. 2015. ISSN 0737-4038.

Wheeler, L. C. and Smith, S. D. Computational Modeling of Anthocyanin Pathway Evolution: Biases, Hotspots, and Trade-offs. *Integrative and Comparative Biology*, 59(3):585–598, 05 2019. ISSN 1540-7063. doi: 10.1093/icb/icz049.

Zufall, R. A. and Rausher, M. D. Genetic changes associated with floral adaptation restrict future evolutionary potential. *Nature*, 428(6985):847–850, Apr. 2004. ISSN 1476-4687.
